# Long-term hybridization in a karst window reveals the genetic basis of eye loss in cavefish

**DOI:** 10.1101/2024.10.25.620266

**Authors:** Riley Kellermeyer, Luis Espinasa, Chris Seidel, William B. Redwine, Rachel L. Moran, Sylvain Bertho, Emma Y. Roback, Claudia Patricia Ornelas-García, Dana Alegre, Kyle Weaver, Jay Unruh, Benjamin Troutwine, Yongfu Wang, Tathagata Biswas, Masato Yoshizawa, Emily Collins, Jennifer Rutkowski, Jordi Espinasa, Suzanne E. McGaugh, Nicolas Rohner

## Abstract

Eye loss is a hallmark trait of animals inhabiting perpetual darkness, yet the precise genetic variants underlying this evolutionary change remain largely unknown. The Mexican tetra (*Astyanax mexicanus*) provides a powerful model for dissecting the genetic basis of eye degeneration, as sighted surface fish and multiple independently evolved blind cave populations remain interfertile; yet despite decades of research and numerous QTL studies, the genetic basis of eye loss has remained unresolved at the level of specific variants. Here, we exploit a rare natural experiment in the Caballo Moro cave, where the collapse of a karst window created a partially illuminated pool inhabited by both fully eyed and completely eyeless cavefish of closely related genetic background. Whole-genome sequencing reveals a long-standing hybrid population between cave and surface lineages, enabling a dramatic refinement of the genetic architecture of eye degeneration to 203 candidate SNPs across 41 genes. Among these, we identified a nonsynonymous mutation in the lens gap-junction protein Connexin-50 (Cx50). CRISPR-based disruption of *cx50* induces early eye loss in surface fish, and F2 laboratory crosses confirm genetic linkage between *cx50* variants and eye size. Additional Cx50 mutations are present in independent cavefish populations and correlate with reduced eye size. Notably, variants in conserved regions of Cx50 also occur in other cave-dwelling fish and subterranean mammals, suggesting repeated evolutionary targeting of this gene. Introduction of the Caballo Moro mutation into mice causes cataracts and reduced eye and lens size, confirming its functional impact. Together, these findings identify the first SNP directly implicated in cavefish eye loss, demonstrate the power of natural hybrid populations to resolve the genetic basis of complex traits, and reveal Cx50 as a case of molecular convergence in vertebrate eye degeneration.

## Main Text

Cave-dwelling and subterranean organisms across diverse taxa have repeatedly evolved traits adapted to life in perpetual darkness, such as the consistent loss or reduction of eyes. Eye degeneration has been hypothesized to result from multiple, non-mutually exclusive processes, including the energetic cost of maintaining visual systems, the relaxation of purifying selection on vision, and indirect selection for other traits that are genetically or developmentally coupled to eye reduction; however, the precise genetic bases of this trait remain poorly understood (*1–6*).

Despite decades of research across a variety of subterranean species, surprisingly little is known about the precise genetic basis of eye loss. Much of the work in this field has focused on *Astyanax mexicanus*, a powerful model for studying cave adaptation. This species exists in two interfertile morphs: sighted surface fish and blind cavefish (*7*). To date, 35 cave populations of *Astyanax* with reduced or absent eyes have been documented (*8–10*). While these do not all represent independent origins, cave-morphs have evolved at least twice independently in the Sierra de Guatemala and Sierra de El Abra regions of Mexico over short evolutionary timescales (*7–15*). As such, eye loss in *A. mexicanus* provides a powerful natural system to investigate repeated evolutionary change.

Quantitative Trait Locus (QTL) analyses in laboratory cave x surface F2 hybrids have linked eye size to at least 14 genomic regions (*6*, *16–22*). However, the resolution of these studies is limited, leaving thousands of potential causal variants within broad genomic intervals. While few cis-regulatory changes have been implicated in eyed degeneration, such as *rx3* and *cystathionine ß-synthase a* (*csba*) (*23–25*), to our knowledge, no causal nucleotide variant has been identified as contributing to eye loss (*4*).

Here, we leveraged a natural experiment in the Caballo Moro cave to identify genes involved in eye degeneration. Located within the Sierra de Guatemala cave system (Fig. 1a, Fig. S1), Caballo Moro formed after a recent ceiling collapse that created a karst window, exposing part of an ∼80 m long cave lake to sunlight (*7*). This results in adjacent illuminated and dark habitats without geographical barriers (Fig. 1b, Fig. S1). Fish inhabiting the illuminated region exhibit surface-like morphology with fully developed and functional eyes (Fig. 1c, d, Fig. S2a), whereas blind cavefish occupy the darker regions (Fig. 1e; *25*). As a result, eyed and eyeless morphs coexist in the same pool, providing a rare opportunity to study the genetic basis of trait divergence in the absence of geographic isolation.

**Fig. 1:**
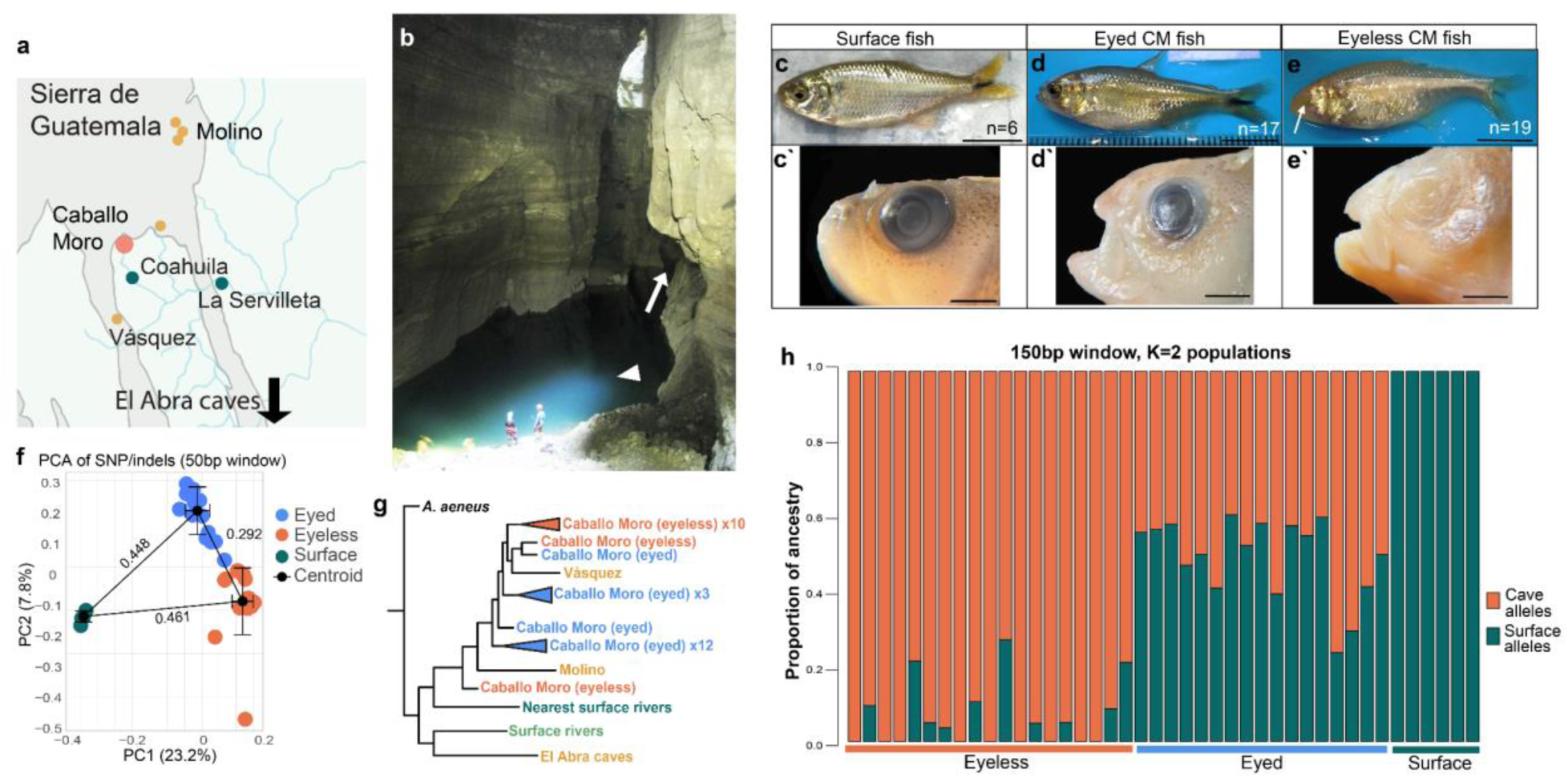
Caballo Moro cavefish and their ancestry. **(a)** Map of the Sierra de Guatemala area caves. Location of Caballo Moro (large orange dot) and other caves (small orange dots) in the Guatemala region. Nearby surface fish were sampled from Coahuila and La Servilleta rivers for comparison (green dots). Arrow indicates that the El Abra caves are further south of the figure (see Fig. S1). **(b)** The Caballo Moro cave pool. Collapse of the roof of the cave exposed part of the cave pool to sunlight (arrowhead) while most of the cave remains in darkness (arrow). **(c-e’)** Examples of fish sampled and sequenced. Whole fish scale is 10mm, eye closeup is 2mm. **(c,c’)** Fish collected from nearby surface rivers. **(d,d’)** Eyed Caballo Moro cavefish collected from the illuminated side of the cave. **(e,e’)** Eyeless Caballo Moro cavefish collected from the dark side of the cave. **(f)** Principal component analysis of SNPs and indels compared to the surface reference genome. Centroids were calculated by phenotype and measured by Euclidean distance. Error bars indicate standard error. **(g)** Consolidated phylogeny of the Caballo Moro cavefish in relation to nearby caves and surface rivers sampled from (*13*). See Fig. S4 for expanded phylogeny. **(h)** Admixture analysis of 679,525 biallelic, statistically unlinked sites. Sites from each individual are sorted into ancestral populations (K=2) with 150bp windows and plotted as admixture coefficients.

Despite the opportunity for gene flow, fish in Caballo Moro are either fully eyed or completely eyeless, with only two individuals exhibiting intermediate eye phenotypes among 75 surveyed fish across this and a previous study (*26*). The striking dichotomy in eye morphology suggests that selective pressures maintain distinct phenotypes despite the close proximity and interfertility of the morphs. Previous genetic analysis suggested a complex evolutionary history for this population, including evidence of hybridization between surface and cave lineages (*26*), providing an opportunity for much higher-resolution genetic mapping of eye loss than is possible in traditional QTL studies.

Through whole genome sequencing of eyed and eyeless Caballo Moro fish, we found that the eyed morph originated from a surface fish invasion of the cave approximately 3,300-4,300 generations ago. Hybridization between these surface-derived fish and the long-established cavefish allowed us to identify relatively few variants differentiating the eyed and non-eyed fish, suggesting these SNPs may be related to eye loss. Using genome-wide association and genotype filtering, we identified a protein-coding mutation in a phylogenetically conserved region of the lens gap junction protein *connexin-50* (*cx50*; also known as *gap junction protein alpha 8b* or lens membrane protein 70). Notably, mutations in this region of CX50 are known in humans to cause cataracts and microphthalmia, highlighting its functional importance for lens development (*27*, *28*). Knocking down *cx50* in surface fish resulted in severe eye degeneration during development. We further identified additional mutations in highly conserved regions of *connexin-50* across independently derived *A. mexicanus* cave populations as well as in other subterranean fishes and mammals. Protein structural modeling of these variants, together with functional validation of a cave-associated mutation in a mouse model, suggests that repeated mutations in *cx50* may represent a case of gene reuse in convergent vertebrate eye degeneration.

### Eyed Caballo Moro cavefish represent longstanding cave-surface hybrids

To determine the origin of the eyed fish from the Caballo Moro (Caballo Moro) cave, we collected 17 light-dwelling eyed fish and 19 dark-dwelling eyeless fish from the cave, along with six surface fish from nearby surface environments (Fig. 1a-e). Anatomical and behavioral characterizations revealed that the eyed Caballo Moro fish, despite having functional eyes, exhibit phenotypes that are typically associated with cavefish, including altered opercular bone orientation and the presence of extra maxillary teeth (Fig. S2b-d; *29*). In two visits 20 years apart, eyed fish were observed to chase eyeless fish from the illuminated side of the pool (*26*). This observation led us to investigate whether behavior contributes to the spatial segregation of the eyed and blind cavefish within the same cave environment. In a controlled resident-intruder assay conducted in the field, we exposed eyed Caballo Moro fish to both surface and eyeless intruders, separated by a glass barrier, and quantified aggressive strikes toward the intruders (Fig. S2e-f; *30*). Eyed Caballo Moro fish attacked both eyeless fish and surface fish, whereas surface fish preferentially attacked other surface fish (p<0.0001, Kruskal-Wallis test). These behavioral findings not only confirm that the eyed Caballo Moro fish have functional vision but also suggest aggression as a potential source of the geographical separation of the different morphs within the cave.

To resolve the population structure and ancestry of the Caballo Moro cavefish, we sequenced all 36 eyed and eyeless individuals from the cave, along with six nearby surface fish, to an average coverage of 8.7X and performed population structure and admixture analysis (Fig. S3a; see Methods). In a principal component analysis (PCA) of SNPs and indels, the eyed and eyeless Caballo Moro fish cluster closely but separate by phenotype (Fig. 1f). Based on Euclidean distances in PCA space, both groups form clusters distinct from surface fish, with the eyed individuals positioned slightly closer to the surface cluster than the eyeless fish (Fig. 1f). Unsupervised k-means clustering similarly identified three distinct groups corresponding to eyed cavefish, eyeless cavefish, and surface fish, with a single eyed individual clustering with the eyeless group (Fig. S3b-c). This population clustering remained consistent when individuals from the nearby Molino cave population were included in the analysis (Fig. S3d-f).

Next, we examined a phylogeny generated by Moran et al. (2023) that includes individuals from the Caballo Moro cave and 26 additional populations (*13*). In this phylogeny, both eyed and eyeless Caballo Moro fish cluster with Vásquez and Molino, two geographically nearby Guatemala cave populations, rather than with surface fish (Fig. 1g, Fig. S4). Apart from two outlier individuals, the eyeless Caballo Moro fish form a well-supported monophyletic clade. In contrast, the eyed Caballo Moro fish are paraphyletic and show weak branch support, suggesting that their evolutionary relationship to the eyeless morph cannot be fully captured by a simple bifurcating tree.

To further investigate the ancestry of the eyed cavefish, we performed admixture analyses across a range of potential ancestral populations (K=1-5). The best supported model identified surface fish and eyeless cavefish as the two parental populations, with the eyed Caballo Moro fish forming an admixed population derived from hybridization between cave and surface lineages (Fig. 1h, Fig. S5a-b). Repeating the analysis with individuals from the nearby Molino cave yielded a similar pattern, with the eyed Caballo Moro fish again showing ancestry consistent with a hybrid population (Fig. S5c-d). This admixture signal provides a potential explanation for the lack of monophyletic clustering of the eyed fish in the phylogenetic analysis.

Additional support for the hybrid origin of the eyed Caballo Moro fish comes from patterns of sequence diversity across populations. Genome-wide nucleotide diversity (π) is approximately 1.5 times higher in surface fish (π = 0.00772 ± 0.0034) than in eyeless cavefish (π = 0.00461 ± 0.0023). Nucleotide diversity in the eyed Caballo Moro fish is intermediate between these groups (π = 0.00567 ± 0.0026), consistent with a hybrid population containing ancestry from both surface and eyeless cave lineages (Table S1).

We further investigated the demographic history of the Caballo Moro populations using coalescent based demographic modeling of derived allele site frequency spectra implemented in fastsimcoal2 (*31*). We tested seven alternative demographic topologies that differed in their patterns of gene flow among populations (Fig. S6a). Models without gene flow received essentially no support (Akaike weight = 0.00), further suggesting that there is gene flow between surface and cave populations.

The best-fitting model among the seven tested topologies (Tree 7; Fig. S6a) suggests that the ancestry of the eyed cavefish traces through an unsampled “ghost” population that is either extinct or has not yet been sampled. In this model, the deepest divergence occurs between this ghost lineage and the sampled surface population approximately 979k generations ago. The eyeless cavefish lineage diverged from the sampled surface lineage roughly 128k generations ago, corresponding to the origin of the cavefish in Caballo Moro cave. Approximately 4300 generations ago, fish from the ghost surface population are inferred to have invaded the cave (Fig. S6b). Following this invasion, the model indicates that the eyed cavefish population derived from this surface lineage and subsequently experienced gene flow with the resident eyeless cave population. Both cave populations also experienced gene flow with the sampled surface population. Notably, inferred gene flow from surface fish into the eyed population is an order of magnitude greater than into the eyeless population (3.88E-04 vs 3.87E-05). Together, these results suggest that the Caballo Moro cave harbors two distinct but interacting lineages: an ancient eyeless cavefish population and eyed cave-surface hybrids derived from a more recent surface invasion.

### Identification of candidate variants underlying eye degeneration

Eye degeneration in *A. mexicanus* is a complex, polygenic trait (*4*, *20*). To identify genetic variants associated with the eyeless phenotype, we employed a cross-sectional genomic approach comparing eyeless, eyed, and surface fish (Fig. S7).

First, we conducted a genome-wide association study (GWAS) to identify genomic regions linked to the eye phenotype. Using a Cochran-Armitage trend test on SNPs and indels in eyeless Caballo Moro fish relative to both eyed and surface fish, we identified seven genomic regions significantly associated with the eyeless phenotype (Cochran-Armitage trend test filtered to -log10(p)>10; Fig. 2a, Table S2). Together, these regions span 467 genes distributed across seven chromosomes.

**Fig. 2:**
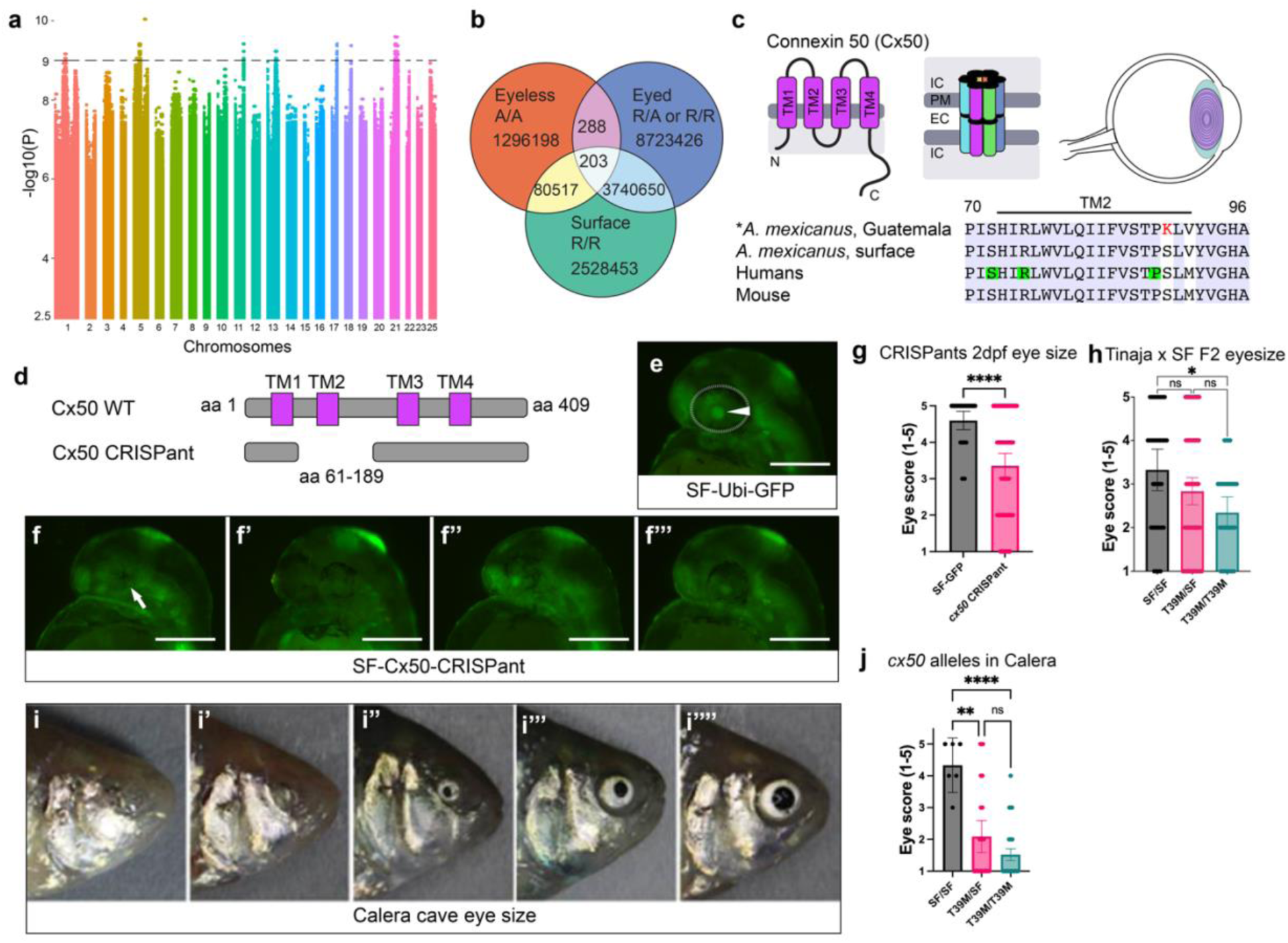
Candidate variant selection and Cx50 CRISPants. **(a)** Manhattan plot of a Cochran-Armitage trend test GWAS comparison of eyeless SNPs to both eyed and surface SNPs. Horizontal line indicates -log10(p)>9. Seven locations associated with eye loss were identified encompassing 7,373 SNPs and 467 genes (Table S2). **(b)** Intersect of SNPs in surface, eyed, and eyeless fish. Surface fish were sorted for having Surface-Reference only alleles (R/R). Eyed fish were filtered to SNPs with at least one reference allele (R/R or R/A). Eyeless fish were filtered to be homozygous for an alternate allele (A/A) compared to the surface reference. 203 SNPs were found that meet these criteria, falling within 40 annotated genes and *cis*-regulatory regions (Table S3). **(c)** Diagram of CX50 and comparison of protein sequence in Guatemala-region caves (including Caballo Moro), surface fish, humans, and mice. Gap junctions are formed by six gap-junction proteins, each with 4 transmembrane domains. Cx50 is expressed primarily in the lens (purple) to connect cells for lens fluidity. Green highlights indicate sites of known human CX50 SNPs that contribute to congenital cataracts or small eyes. Red text indicates the site modified in eyeless cavefish when compared against eyed Caballo Moro and surface fish **(d)** Schematic of Cx50 TM2 region targeted in CRISPants. CRISPants were defined as more than 99% of sequenced reads with a deletion of the target sequence. **(e-f)** 2dpf surface fish with transgenic ubiquitin-GFP (SF-Ubi-GFP; *34*) expression from the same clutch. Ubi-GFP allows better visualization of the lens (arrowhead). Bar is 500μm. **(e)** Un-injected SF-Ubi-GFP with wild-type eye size, a score of 5 (n=15 fish; both eyes scored, n=30). Arrowhead indicates the lens; outline is the eye. **(f-f”’)** The spectrum of eye phenotypes in Cx50 CRISPants ranked by increasing eye size, representing eye size scores from 1 through 4 from left to right (n=37 fish; both eyes scored, n=74). Note the complete collapse of eye and pigment cells into a single point in score 1 (arrow), and the absence of a lens in score 2. A lens is generally visualized in scores above 3. **(g)** Quantification of eye size score between uninjected transgenic fish and CRISPants (p<0.0001, two-tailed Mann-Whitney U test, error bars are 95% confidence interval). **(h)** Quantification of adult Tinaja and surface fish F2 hybrid eye sizes by *cx50* genotype. Homozygous F2 hybrids (n=16, both eyes measured) have smaller eyes than surface fish (n=17, both eyes measured; p=0.0163, Kruskal-Wallis test). N=45 heterozygous fish, both eyes measured. **(i-i’’’’)** Sample of eye sizes from the hybrid population in the Calera cave, with scores of 1-5 (n=119). **(j)** Quantification of eye scores in Calera cavefish by Cx50-T39M genotypes. Surface fish allele (SF) homozygous (SF/SF) versus heterozygous (T39M/SF) p=0.0023; T39M/SF versus homozygous cave allele (T39M/T39M) p=0.2065; Surface homozygous versus cave homozygous (T39M/T39M) **** p<0.0001; Kruskal-Wallis test with Dunn’s multiple comparison, error bars are 95% confidence interval.

Second, we identified eyeless-specific variants by filtering for SNPs that were homozygous for the alternate allele in all eyeless individuals while retaining the reference allele in both eyed cavefish and surface fish. We then intersected these variants with loci carrying the reference allele in both the eyed cavefish and surface fish. This filtering step yielded 203 candidate SNPs (Fig. 2b, Table S3). The vast majority of these variants (199/203; 98%) occur in non-coding regions of the genome, consistent with previous estimates that much of the genetic basis of morphological evolution resides in regulatory regions (*33*).

Finally, we developed a custom database to further prioritize candidates (Table S4). We compiled a list of genes that have experienced hard selective sweeps in cavefish from Moran et al., (2023), indicating positive selection (*13*). We also compiled a list of genes that fall within eye phenotypes in large-scale F2 cave-surface hybrid QTL studies (*20*). The last component of our database is a list of zebrafish genes that have eye-specific gene ontology. We intersected our list of eyeless Caballo Moro candidate genes with this custom database and found that variants in or near five genes meet the criteria of all datasets - *cx50*, *apcdd1l*, *agrn*, *rpgrip1*, and *tubgcp3* (Fig. S7).

To prioritize candidate genes, we evaluated coding variation and predicted functional impact among the variants in these five candidate genes. Among them, *cx50* was the only gene carrying a nonsynonymous, high impact protein coding mutation, highlighting it as a strong candidate for functional involvement in eye degeneration. In eyeless Caballo Moro fish, *connexin-50* (*cx50*) contains a serine to lysine substitution at position 89 (S89K). This variant stood out for several reasons. First, the closest human homolog, CX50 (or GJA8), encodes a lens-specific gap-junction protein essential for lens development, homeostasis, and transparency (*34*). Second, the S89K substitution affects a highly conserved residue within a transmembrane domain critical for channel function. Third, *cx50* lies within a genomic region showing a strong signature of hard selective sweep in cavefish populations, consistent with positive selection (*13*). Finally, this locus falls within previously identified eye-associated QTL intervals, further supporting a role in the evolution of the eyeless phenotype (*6*, *20–22*).

In teleosts, *cx50* has a paralog on a different chromosome, *gja8a*, which carries a single amino acid insertion in eyeless Caballo Moro fish and poor sequence similarity to mammalian CX50 (Fig. S8a, *35*). However, this locus does not show signatures of selection and does not fall within eye-size QTL intervals (*13*, *20*). Interestingly, larger deletions in *gja8a* occur in other cavefish species, suggesting that additional connexin-family variation may contribute to eye evolution in cave environments.

### Connexin-50 and eye degeneration in *A. mexicanus*

In *A. mexicanus*, the lens acts as a master regulator of eye regression, as reciprocal lens transplantation experiments between cave and surface fish demonstrated that the cavefish lens is sufficient to drive eye degeneration (*36*). Both morphotypes initiate eye development similarly until approximately 24 hours post fertilization (hpf), after which the cavefish lens undergoes extensive apoptosis that ultimately leads to degeneration of the entire eye (*37*, *38*).

In mammals, *CX50* is primarily expressed in the lens, where it forms intercellular gap junction channels composed of 12 connexin subunits that connect adjacent cells (Fig. 2c; *28*, *39*). CX50 is the only connexin present in interior lens fibers and is essential for maintaining lens homeostasis and transparency (*34*, *40*). In humans, more than 40 disease-associated mutations in *CX50* disrupt gap junction function and cause congenital cataracts and microphthalmia (reduced eye size) through impaired lens fiber differentiation and increased apoptosis *47*, *48*). Notably, four of these pathogenic variants occur immediately adjacent to the S89K mutation identified in eyeless Caballo Moro fish (*43–49*).

To evaluate the role of Cx50 in *A. mexicanus* eye development, we first confirmed that the gene is expressed in the developing lens. RNA-seq of lenses isolated from surface fish at 24 hours post fertilization (hpf) detected robust *cx50* expression during this critical stage of eye development. The *cx50* gene in surface fish lenses had a mean transcripts per million (TPM) value of 374 ± 75, which is above the 98th percentile of TPM values for genes detected in the experiment. To test whether Cx50 is required for eye formation, we generated surface fish F0 CRISPants using *cx50*-targeted CRISPR guides (Fig. 2d-g, Fig. S8b). Disruption of *cx50* resulted in severe eye defects: mutants displayed either small or no eyes by 2 days post-fertilization (dpf) (Fig. 2g, p<0.0001, Mann-Whitney U test). Additionally, ∼5% of *cx50* CRISPants displayed a cyclops phenotype (Fig. S9). By 6 dpf, *cx50* mutants displayed a spectrum of eye phenotypes similar to those observed at 2dpf, although completely eyeless individuals were no longer observed (Fig. S9a-e). We suspect that the initial formation of a lens is required for proper development, as no 6dpf fish completely lacked a lens, in comparison to 2dpf fish (Fig. S9d), though the lenses of 6dpf CRISPants showed a qualitative increase in apoptosis (Fig. S10). These results demonstrate that Cx50 plays a critical role in lens-dependent eye development in *A. mexicanus* which is in line with its essential role in early postnatal lens growth and cell division in mice (*50–52*).

### Independent mutations in *cx50* across *A. mexicanus* cavefish lineages

*Astyanax mexicanus* cavefish evolved through at least two independent colonization events from ancestral surface populations invaded caves in the Sierra de Guatemala and the Sierra de El Abra regions (*11–13*, *53*). Given the close relationship between Caballo Moro cavefish and other Guatemala populations, we sequenced *cx50* in additional caves from this region, including Molino, Vásquez, Escondido, and Jineo. All populations shared the same S89K allele identified in Caballo Moro.

We next examined whether the same mutation occurs in cavefish from the independently derived Sierra de El Abra lineage. We used publicly available sequences of Pachón, Los Sabinos, Tinaja, Curva, and Calera populations and found that these cavefish retain the ancestral residue at position S89. However, all El Abra populations carry a different mutation in *cx50*: a threonine to methionine substitution at position 39 (T39M). This residue lies within the first transmembrane domain of CX50 and corresponds to a position where human mutations (e.g., T39R) are known to cause congenital cataracts (*28*, *54*, *55*). To test whether this mutation is associated with eye size variation, we generated F2 hybrids by crossing surface fish with Tinaja cavefish from the El Abra lineage. We genotyped 78 F2 hybrids for *cx50* and measured eye size. We found a correlation between reduced eye size and the presence of the Cx50-T39M variant (Fig. 2h, p=0.0163, Kruskal-Wallis test).

To evaluate the effects of this mutation under natural environmental conditions, we conducted field studies in the Calera cave, which is part of the El Abra cave lineage (*9*). Fish from this population exhibit a continuous spectrum of eye sizes similar to that observed in F2 hybrids (*9*). We scored eye sizes for 119 wild-caught Calera individuals and correlated these measurements with their *cx50* genotypes. The T39M allele showed a strong association with reduced eye size in this natural population (p<0.0001, Kruskal-Wallis test; Fig. 2i-j), demonstrating a robust genetic link between *cx50* variation and eye reduction across distinct cavefish populations in the wild.

Together, these results indicate that mutations in *cx50* contribute to eye size variation in *A. mexicanus*. Disruption of *cx50* in surface fish produces severe eye defects, while distinct *cx50* mutations are associated with reduced eye size in independently evolved cavefish lineages. Moreover, the *cx50* locus lies within genomic regions showing signatures of positive selection in both the Guatemala and El Abra cave lineages. These findings suggest that selection has repeatedly targeted *cx50* during the evolution of eye degeneration, even in populations experiencing ongoing gene flow with surface fish.

### Variants in Cx50 across other cavefishes and subterranean mammals

Given the independent mutations in *cx50* identified in two *A. mexicanus* cavefish lineages, we next investigated whether variation in this gene occurs in other cave-adapted fishes. We analyzed publicly available sequences from 19 cavefish species and their closest available surface relatives with a particular focus on the first 245 N-terminal amino acids of Cx50 which encompass the N-terminal helix and the first two transmembrane domains (Figure 3a, Fig. S11, Table S5). These regions are highly conserved across fish, showing approximately 98% amino acid sequence homology among the species examined. Notably, eight of the 19 cavefish species carry non-synonymous substitutions in regions of *cx50* known to have phenotypic effects in humans, such as the N-terminal helix (NTH) and first two transmembrane domains (TM1, TM2, Fig. S11). Notably, one of the other cavefish analyzed, the Chinese blind golden-line barbel, *Sinocyclocheilus anophthalmus*, has an S89F mutation at the same residue as the Guatemala lineage fish and a R33Q mutation at the same residue as a known human cataract phenotype (Fig. S11; *62*, *63*).

**Fig. 3:**
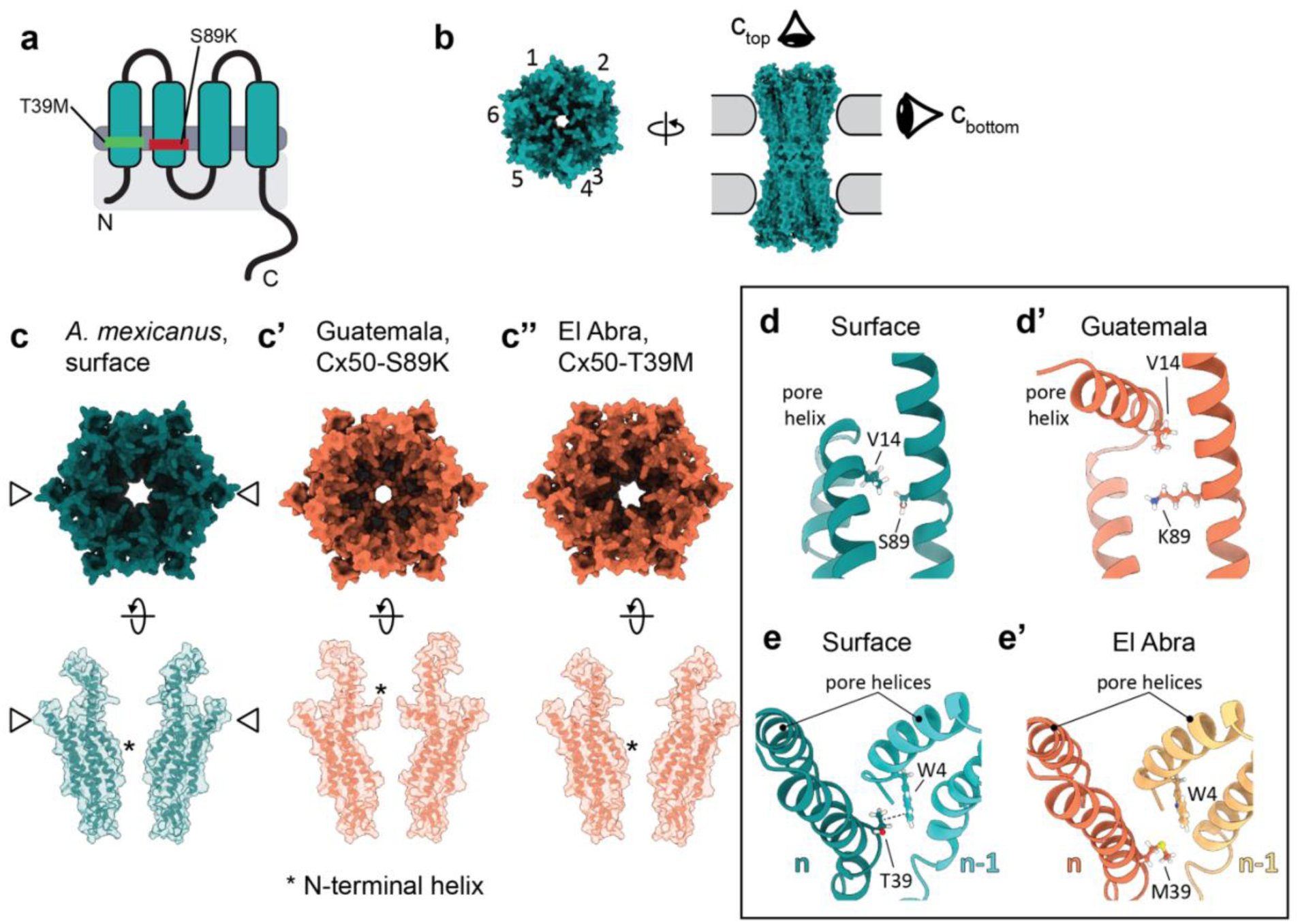
Predicted Cx50 structure in *Astyanax mexicanus*. **(a)** Connexin 50 is a transmembrane protein with four domains and a disorganized intercellular C-terminus. In *A. mexicanus* cave populations, T39M in El Abra lineage cavefish is in transmembrane 1 (TM1), S89K in Guatemala lineage cavefish is in transmembrane 2 (TM2). **(b)** CX50 modeling of gap junctions across the plasma membrane. Six CX50 proteins form a hemichannel in one cell membrane and couples with a hemichannel on an adjacent cell. The N-terminal helix (NTH) in the interior of the connexin pore contributes to ion selectivity and voltage gating and controls the stability of open and closed states by interacting with transmembranes 1 and 2. **(c-c’’)** Modeling of the interior of the pore in *A. mexicanus* variants. Top figures are models visualized down the center of the pore. Bottom figures are cross sections of one side of the pore between subunits 6 and 3. Positions of the NTH are indicated with an asterisk. Note that the Guatemala allele results in the N-terminus helix flipped up into the pore. **(d-e)** Molecular renderings of the altered variants in cavefish and the key amino acids with which they interact. **(d)** Comparison of the interaction between aa89 in TM2 and the N-terminal helix of a single Cx50. In the surface protein, S89 and V14 pack closely together. **(d’)** In the S89K variant of Guatemala lineage cavefish, lysine increases steric hinderance and alters the hydrophobic interaction with V14, pushing the NTH into the pore. **(e)** Comparison of the interaction between aa39 (n) and W4 in the N-terminal helix of an adjacent Connexin-50 protein (n-1). M39 reduces the stability of the NTH across the subunits and increases packing away from the pore.

We extended this analysis to subterranean mammals, which have also evolved convergent eye degeneration (*58–60*). Teleost Cx50 and mammalian CX50 share approximately 83% amino acid sequence similarity in the first 245 amino acids, highlighting the high conservation of this protein across vertebrates. We therefore compared *Cx50* sequences from five subterranean-adapted mammal genomes and their closest available surface relatives (Fig. S12, Table S5). Four species, the Upper Galilee Mountains blind mole-rat (*Nannospalax galili*), Damara mole-rat (*Fukomys damarensis*), Iberian mole (*Talpa occidentalis*), and star-nosed mole (*Condylura cristata*) harbor subterranean specific, nonsynonymous substitutions within the NTH or the first two transmembrane domains.

To evaluate the potential functional effects of these substitutions, we performed *in silico* structural analyses of Cx50/CX50 sequences from seven cavefish species, three mammals, and their closest surface relatives. We used AlphaFold2, together with the cryo-electron microscopy-resolved CX50 structure, to model the intracellular channel conformation of the N-terminal 245 residues of Cx50/CX50 (Fig. 3b; *45*, *67*, *68*). Structural models were visualized in ChimeraX to examine potential effects on gap junction architecture (*63*). These static models allow for the assessment of gross changes to the pore, such as the open-state conformation, orientation of functional elements like the N-terminal helix, and electrostatic potential (Fig. 3). Previous studies have shown that mutations within these elements of the protein, especially in the inner pore regions of the NTH, TM1, and TM2, can drastically alter the selectivity and conductance of the pore. Disruption of the CX50 pore is known to impair antioxidant and growth factor distribution, leading to cataracts, reduced cellular proliferation, and increased apoptosis (*64–66*). We therefore examined whether the cave-associated mutations alter these structural features of the CX50 pore.

We first modeled the *Astyanax* Cx50 variants found in our cave populations – S89K and T39M (Fig. 3c-e’). In the cryo-electron microscopy structure of CX50, hydrophobic interactions between the NTH and TM2 are required for proper voltage gating and open channel stability (*67*). Structural modeling of the S89K variant found in Caballo Moro and other Guatemala lineage cavefish predicts that the NTH is repositioned into the channel opening due to increased steric hinderance of K89 and the hydrophobic side of the NTH (Fig. 3d-d’, Fig. S13). This repositioning results in a stabilized closed-state conformation and a smaller predicted interior pore that has the potential to disrupt channel function (Fig. S13b). Similar to the Caballo Moro Cx50 proteins, the two mutations in the Chinese blind golden-line barbel *S. anophthalmus* (S89F, R33Q) are predicted to reposition the NTH toward the pore, potentially constricting the channel similar to the Guatemala variants (Fig. S14a-c).

In contrast, the T39M mutation carried by El Abra cavefish is predicted to increase pore diameter (Fig. 3c”). In previously resolved and tested structures, T39 in TM1 on one subunit interacts with W4 in the NTH of an adjacent subunit to maintain a set distance between the domains (Fig. 3e-e’; *61*). M39 disrupts this interaction, which ultimately increases the predicted pore size in the most intracellular portion of the pore (Fig. 3c, e’, Fig. S13c). This predicted structural effect resembles that of the human T39R mutation which stabilizes the open channel conformation and enhances hemichannel activity. Increased hemichannel conductance results in accumulation of calcium ions and depletion of intracellular metabolites, ultimately triggering apoptosis and cataracts (*55*).

Among the subterranean mammals examined, CX50 from the Damara mole-rat, *F. damarensis*, showed the most remarkable predicted structure change (Fig. S14d-f). The Damara mole-rat has small eyes and heavy eyelids that impair vision (*68*). In this species, CX50 carries a negatively charged glutamic acid change to a polar, uncharged glutamine in the intracellular N-terminal helix (E12Q). In several AlphaFold models, the E12Q substitution resulted in a repositioning of the NTH into the pore and a narrowing of the pore diameter relative to a mouse structural model (Fig. S14d-e). In models where the NTH retained the canonical downward conformation, the mutation was still predicted to alter pore properties. Because residue E12 faces the interior of the pore, replacing this negatively charged residue is predicted to affect ion conductance, as CX50 has selectivity for cations that depends upon a number of negatively charged pore residues (Fig. S14d-f; *73*). Together, these structural models suggest that Cx50/CX50 variants found in subterranean-dwelling creatures repeatedly affect key pore-forming regions of the channel and represent an example of molecular convergence with the same gene evolving mutations that affect protein function in response to similar environmental pressures.

To assess the validity of our protein structure-based prediction pipeline, we selected the eyeless Caballo Moro Cx50-S89K variant for functional testing *in vivo* (Fig. 4). Using CRISPR-Cas9 genome editing, we introduced the S89K mutation into the endogenous *Cx50* locus in mice. Mice homozygous for the CX50-S89K mutation developed congenital cataracts (Fig. 4c-d). More strikingly, both CX50-S89K heterozygotes and homozygote mice showed significantly reduced eye area and lens area compared to wildtype CX50 mice (Fig. 4a, b, e-i, multiple linear regression controlling for age). The significantly reduced eye and lens sizes in CX50-S89K heterozygotes indicates a dominant effect of this allele that has also been reported for certain human SNPs (*43*, *46*, *69*, *70*). These results *in vivo* not only confirm the contribution of the CX50-S89K mutation to eye size but also demonstrate the robustness of our protein modeling pipeline in identifying functionally significant convergent evolutionary changes. Functional validation of the other Cx50 variants in subterranean animals in the future would be valuable for further understanding the role of this lens protein in evolution.

**Fig. 4:**
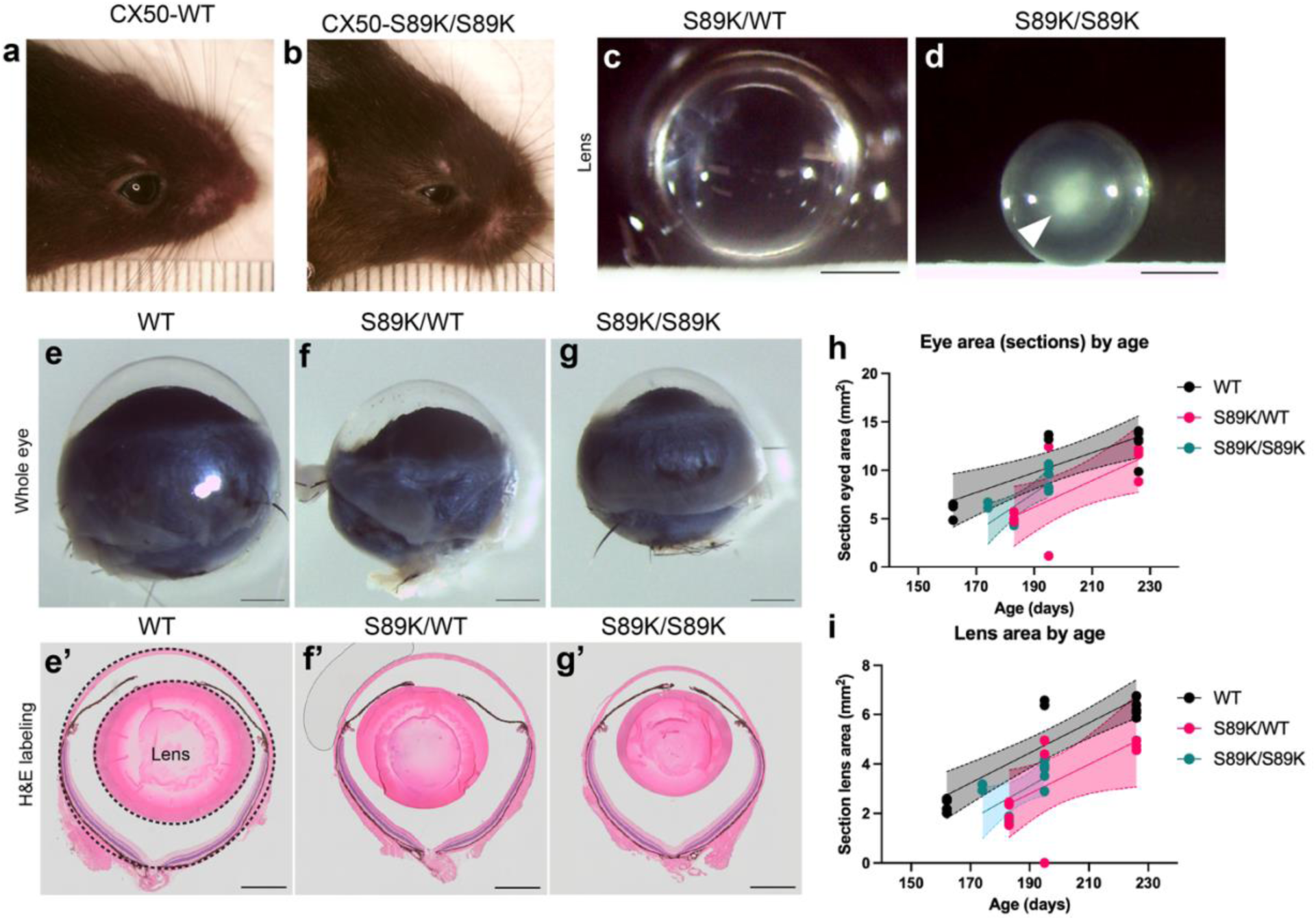
Testing SNPs with predicted CX50 protein function *in vivo*. (a-b) Differences in eye size between wild-type and homozygous CX50-S89K mice can be visualized externally. Ruler is in mm. **(c-d)** Imaging of dissected lens from adult heterozygous and homozygous CX50-S89K littermates (n=9). **(c)** Heterozygous lenses are transparent. **(d)** CX50-S89K mutant lenses are opaque with a cataract (arrowhead). **(e-g’)** Eye and lens visualization. Bar is 1mm. **(e-g)** Whole eye visualization of WT, heterozygous, and homozygous for the CX50-S89K allele in adult mice. **(e’-g’)** H&E labeling of section of the eye for measurements of the eye and lens. Outlines indicate the diameter of the eye and lens. **(h-i)** Quantification of eye area from H&E labels using multiple linear regression with age and genotype as predictors (Eye area ∼ Intercept + Genotype + Age. 3-5 sections were averaged for each eye (n=18 mice). **(h)** Eye area was analyzed using multiple linear regression with age and genotype as predictors. After accounting for age, eye area was smaller for CX50-S89K heterozygotes and homozygotes (multiple linear regression, overall model F(3,28)=16.33, p<0.0001, R^2^=0.64; CX50-S89K/WT: β=-3.17, p=0.0045; CX50-S89K/S89K: β=-2.14, p=0.0448). **(i)** Lens area was analyzed using multiple linear regression with age and genotype as predictors. After accounting for age, lens area was smaller for CX50-S89K heterozygotes and homozygotes (multiple linear regression, overall model F(3,31)=24.22, p<0.0001, R^2^=0.70; CX50-S89K/WT: β=-1.66, p=0.0007; CX50-S89K/S89K: β=-1.36, p=0.0044) compared to wildtype CX50.

In this study, we leveraged a unique *Astyanax mexicanus* cavefish population to identify the first specific genetic variant contributing to eye degeneration in this system. Previous work has identified genomic regions associated with eye size, but these regions often contain thousands to millions of candidate variants. By exploiting a naturally hybridizing cave population, we were able to narrow this space to a small set of candidate variants, including a protein-coding mutation in Connexin50 that we functionally validated *in vivo*. Although eye loss in *A. mexicanus* is a complex trait shaped by multiple genetic factors, our results identify *cx50* as a key locus associated with eye reduction across independent cavefish populations. Importantly, the remaining ∼200 candidate variants identified here provide a tractable set of loci that may enable the genetic architecture of cavefish eye degeneration to be resolved in future work.

Beyond *A. mexicanus*, we detected additional Cx50 variants in other cave-adapted fishes and subterranean mammals that are predicted to alter gap junction function. Because *Cx50* is expressed almost exclusively in the lens, mutations in this gene are less likely to affect other tissues, potentially reducing pleiotropic constraints. This restricted expression may make cx50 particularly susceptible to repeated evolutionary modification. Together, these findings suggest that locus reuse in cx50 may represent a recurrent molecular pathway underlying eye degeneration across subterranean vertebrates.

## Acknowledgements

We thank M. Kirkman, J. Yonke, K. Zueckert-Gaudenz, and A. Kenzior for sample preparation and processing. We additionally thank students of the BIOL320 Genetics courses at Marist University for DNA sequencing and morphology measurements of Tinaja and Calera cavefish. We are grateful for N. Morris and M. Frangello for mouse care and lens imaging; the Stowers Aquatics facility for fish care and support; J. Hunter for eye quantifications; and J. VanCampen, G. Kong, and J.K. Medley for computational assistance. We thank three anonymous reviewers for their careful, rigorous, and constructive feedback, which greatly improved the clarity and quality of this manuscript.

## Funding

Stowers Institute for Medical Research (RK, SB, DA, NR)

American Association of University Women Postdoctoral Research Leave Fellowship (RK)

National Institute of Health grant R24OD030214 (NR)

National Science Foundation grant DEB 2316783 (RK, EYR, SEM) Programa de Apoyo a Proyectos de Investigación e Innovación Tecnológica (PAPIIT) UNAM, IN214624 (COPG)

Marist University (LE, EC, JE, JR)

## Author contributions

Conceptualization: LE, NR, RK, EYR, SEM Writing and editing: RK, NR, SEM, EYR Fieldwork: CPOG, LE, NR, EC, JE, JR

Cavefish phenotyping: LE, JE RK Sequencing: KW, LE

Computational analysis: RK, CS, RLM, SEM, EYR, DA Cavefish CRISPants: SB BT

Protein modeling: WBR JU Mouse phenotyping: YW.

## Data, code, and materials availability

All genome sequencing data generated for this project is available on NCBI under PRJNA841601, SRA accessions are listed in Table S6. RNA sequencing data is available with GEO accession number GSE317165. The surface fish *Astyanax mexicanus* reference genome (AstMex3_surface Accession # GCF_023375975.1) is available on NCBI. Custom code for data processing and analysis is available on GitHub (https://github.com/rikellermeyer/Caballo_Moro).

## List of Supplementary Materials

Materials and Methods

Figs. S1 to S14, Table S5

Tables S1-S4, S6

Data S1 References

## References

1. K. Hüppop, Oxygen consumption of Astyanax fasciatus (Characidae, Pisces): a comparison of epigean and hypogean populations. Environ. Biol. Fishes 17, 299–308 (1986).

2. H. Wilkens, “Evolution and Genetics of Epigean and Cave Astyanax fasciatus (Characidae, Pisces)” in Evolutionary Biology (1988)vol. 23, pp. 271–367.

3. D. Moran, R. Softley, E. J. Warrant, The energetic cost of vision and the evolution of eyeless Mexican cavefish. Sci. Adv. 1, e1500363 (2015).

4. I. Sifuentes-Romero, A. M. Aviles, J. L. Carter, A. Chan-Pong, A. Clarke, P. Crotty, D. Engstrom, P. Meka, A. Perez, R. Perez, C. Phelan, T. Sharrard, M. I. Smirnova, A. J. Wade, J. E. Kowalko, Trait Loss in Evolution: What Cavefish Have Taught Us about Mechanisms Underlying Eye Regression. Integr. Comp. Biol. 63, 393–406 (2023).

5. Y. Yamamoto, M. S. Byerly, W. R. Jackman, W. R. Jeffery, Pleiotropic functions of embryonic sonic hedgehog expression link jaw and taste bud amplification with eye loss during cavefish evolution. Dev. Biol. 330, 200–211 (2009).

6. M. Protas, M. Conrad, J. B. Gross, C. Tabin, R. Borowsky, Regressive Evolution in the Mexican Cave Tetra, Astyanax mexicanus. Curr. Biol. 17, 452–454 (2007).

7. R. W. Mitchell, W. H. Russell, W. R. Elliott, Mexican Eyeless Characin Fishes, Genus Astyanax: Environment, Distribution, and Evolution. Spec. Publ. Mus. Tex. Tech Univ. (1977).

8. L. Espinasa, L. Legendre, J. Fumey, M. Blin, S. Rétaux, M. Espinasa, A new cave locality for Astyanax cavefish in Sierra de El Abra, Mexico. Subterr. Biol. 26, 39–53 (2018).

9. L. Espinasa, C. P. Ornelas-García, L. Legendre, S. Rétaux, A. Best, R. Gamboa-Miranda, H. Espinosa-Pérez, P. Sprouse, Discovery of Two New Astyanax Cavefish Localities Leads to Further Understanding of the Species Biogeography. Diversity 12, 368 (2020).

10. R. Miranda-Gamboa, L. Espinasa, M. de los A. Verde-Ramírez, J. Hernández-Lozano, J. L. Lacaille, M. Espinasa, C. P. Ornelas-García, A new cave population of Astyanax mexicanus from Northern Sierra de El Abra, Tamaulipas, Mexico. Subterr. Biol. 45, 95–117 (2023).

11. M. Garduño-Sánchez, J. Hernández-Lozano, R. L. Moran, R. Miranda-Gamboa, J. B. Gross, N. Rohner, W. R. Elliott, J. Miller, L. Lozano-Vilano, S. E. McGaugh, C. P. Ornelas-García, Phylogeographic relationships and morphological evolution between cave and surface Astyanax mexicanus populations (De Filippi 1853) (Actinopterygii, Characidae). Mol. Ecol. 32, 5626–5644 (2023).

12. J. B. Gross, The complex origin of Astyanax cavefish. BMC Evol. Biol. 12, 105 (2012).

13. R. L. Moran, E. J. Richards, C. P. Ornelas-García, J. B. Gross, A. Donny, J. Wiese, A. C. Keene, J. E. Kowalko, N. Rohner, S. E. McGaugh, Selection-driven trait loss in independently evolved cavefish populations. Nat. Commun. 14, 2557 (2023).

14. J. Fumey, H. Hinaux, C. Noirot, C. Thermes, S. Rétaux, D. Casane, Evidence for late Pleistocene origin of Astyanax mexicanus cavefish. BMC Evol. Biol. 18, 43 (2018).

15. M. Policarpo, L. Legendre, I. Germon, P. Lafargeas, L. Espinasa, S. Rétaux, D. Casane, The nature and distribution of putative non-functional alleles suggest only two independent events at the origins of Astyanax mexicanus cavefish populations. BMC Ecol. Evol. 24, 41 (2024).

16. R. Borowsky, H. Wilkens, Mapping a Cave Fish Genome: Polygenic Systems and Regressive Evolution. J. Hered. 93, 19–21 (2002).

17. S. E. McGaugh, J. B. Gross, B. Aken, M. Blin, R. Borowsky, D. Chalopin, H. Hinaux, W. R. Jeffery, A. Keene, L. Ma, P. Minx, D. Murphy, K. E. O’Quin, S. Rétaux, N. Rohner, S. M. J. Searle, B. A. Stahl, C. Tabin, J.-N. Volff, M. Yoshizawa, W. C. Warren, The cavefish genome reveals candidate genes for eye loss. Nat. Commun. 5, 5307 (2014).

18. K. E. O’Quin, M. Yoshizawa, P. Doshi, W. R. Jeffery, Quantitative Genetic Analysis of Retinal Degeneration in the Blind Cavefish Astyanax mexicanus. PLOS ONE 8, e57281 (2013).

19. W. C. Warren, T. E. Boggs, R. Borowsky, B. M. Carlson, E. Ferrufino, J. B. Gross, L. Hillier, Z. Hu, A. C. Keene, A. Kenzior, J. E. Kowalko, C. Tomlinson, M. Kremitzki, M. E. Lemieux, T. Graves-Lindsay, S. E. McGaugh, J. T. Miller, M. T. M. Mommersteeg, R. L. Moran, R. Peuß, E. S. Rice, M. R. Riddle, I. Sifuentes-Romero, B. A. Stanhope, C. J. Tabin, S. Thakur, Y. Yamamoto, N. Rohner, A chromosome-level genome of Astyanax mexicanus surface fish for comparing population-specific genetic differences contributing to trait evolution. Nat. Commun. 12, 1447 (2021).

20. J. Wiese, E. Richards, J. E. Kowalko, S. E. McGaugh, Quantitative trait loci concentrate in specific regions of the Mexican cavefish genome and reveal key candidate genes for cave-associated evolution. J. Hered., esae040 (2025).

21. M. Protas, I. Tabansky, M. Conrad, J. B. Gross, O. Vidal, C. J. Tabin, R. Borowsky, Multi-trait evolution in a cave fish, Astyanax mexicanus. Evol. Dev. 10, 196–209 (2008).

22. A. Lyon, A. K. Powers, J. B. Gross, K. E. O’Quin, Two – three loci control scleral ossicle formation via epistasis in the cavefish Astyanax mexicanus. PLOS ONE 12, e0171061 (2017).

23. J. Leclercq, J. Torres-Paz, M. Policarpo, F. Agnès, S. Rétaux, Evolution of the regulation of developmental gene expression in blind Mexican cavefish. Development 151, dev202610 (2024).

24. L. Ma, A. V. Gore, D. Castranova, J. Shi, M. Ng, K. A. Tomins, C. M. van der Weele, B. M. Weinstein, W. R. Jeffery, A hypomorphic cystathionine ß-synthase gene contributes to cavefish eye loss by disrupting optic vasculature. Nat. Commun. 11, 2772 (2020).

25. D. Shennard, I. Sifuentes-Romero, R. Ambosie, J. Abdelaziz, E. R. Duboue, J. E. Kowalko, The rx3 Gene Contributes to the Evolution of Eye Loss in the Cavefish Astyanax mexicanus. Evol. Dev. 27, e70011 (2025).

26. L. Espinasa, R. Borowsky, EYED CAVE FISH IN A KARST WINDOW. J. Cave Karst Stud. 62, 180–183 (2000).

27. V. M. Berthoud, J. Gao, P. J. Minogue, O. Jara, R. T. Mathias, E. C. Beyer, Connexin Mutants Compromise the Lens Circulation and Cause Cataracts through Biomineralization. Int. J. Mol. Sci. 21, 5822 (2020).

28. F. Ceroni, D. Aguilera-Garcia, N. Chassaing, D. A. Bax, F. Blanco-Kelly, P. Ramos, M. Tarilonte, C. Villaverde, L. R. J. da Silva, M. J. Ballesta-Martínez, M. J. Sanchez-Soler, R. J. Holt, L. Cooper-Charles, J. Bruty, Y. Wallis, D. McMullan, J. Hoffman, D. Bunyan, A. Stewart, H. Stewart, K. Lachlan, A. Fryer, V. McKay, J. Roume, P. Dureau, A. Saggar, M. Griffiths, P. Calvas, C. Ayuso, M. Corton, N. K. Ragge, DDD Study, New GJA8 variants and phenotypes highlight its critical role in a broad spectrum of eye anomalies. Hum. Genet. 138, 1027–1042 (2019).

29. A. D. S. Atukorala, T. A. Franz-Odendaal, Genetic linkage between altered tooth and eye development in lens-ablated *Astyanax mexicanus*. Dev. Biol. 441, 235–241 (2018).

30. Y. Elipot, H. Hinaux, J. Callebert, S. Rétaux, Evolutionary Shift from Fighting to Foraging in Blind Cavefish through Changes in the Serotonin Network. Curr. Biol. 23, 1–10 (2013).

31. L. Excoffier, N. Marchi, D. A. Marques, R. Matthey-Doret, A. Gouy, V. C. Sousa, fastsimcoal2: demographic inference under complex evolutionary scenarios. Bioinformatics 37, 4882–4885 (2021).

32. B. A. Stahl, R. Peuß, B. McDole, A. Kenzior, J. B. Jaggard, K. Gaudenz, J. Krishnan, S. E. McGaugh, E. R. Duboue, A. C. Keene, N. Rohner, Stable transgenesis in Astyanax mexicanus using the Tol2 transposase system. Dev. Dyn. Off. Publ. Am. Assoc. Anat. 248, 679–687 (2019).

33. A. V. Gore, K. A. Tomins, J. Iben, L. Ma, D. Castranova, A. E. Davis, A. Parkhurst, W. R. Jeffery, B. M. Weinstein, An epigenetic mechanism for cavefish eye degeneration. *Nat*. Ecol. Evol. 2, 1155–1160 (2018).

34. V. M. Berthoud, A. Ngezahayo, Focus on lens connexins. BMC Cell Biol. 18, 6 (2017).

35. M. Policarpo, L. Legendre, I. Germon, P. Lafargeas, L. Espinasa, S. Rétaux, D. Casane, The nature and distribution of putative non-functional alleles suggest only two independent events at the origins of Astyanax mexicanus cavefish populations. BMC Ecol. Evol. 24, 41 (2024).

36. Y. Yamamoto, W. R. Jeffery, Central Role for the Lens in Cave Fish Eye Degeneration. Science 289, 631–633 (2000).

37. A. Alunni, A. Menuet, E. Candal, J.-B. Pénigault, W. R. Jeffery, S. Rétaux, Developmental mechanisms for retinal degeneration in the blind cavefish Astyanax mexicanus. J. Comp. Neurol. 505, 221–233 (2007).

38. W. R. Jeffery, D. P. Martasian, Evolution of Eye Regression in the Cavefish Astyanax: Apoptosis and the Pax-6 Gene1. Am. Zool. 38, 685–696 (1998).

39. J. B. Myers, B. G. Haddad, S. E. O’Neill, D. S. Chorev, C. C. Yoshioka, C. V. Robinson, D. M. Zuckerman, S. L. Reichow, Structure of native lens connexin-46/50 intercellular channels by CryoEM. Nature 564, 372–377 (2018).

40. T. W. White, D. A. Goodenough, D. L. Paul, Targeted Ablation of Connexin50 in Mice Results in Microphthalmia and Zonular Pulverulent Cataracts. J. Cell Biol. 143, 815–825 (1998).

41. L. Li, D.-B. Fan, Y.-T. Zhao, Y. Li, Z.-B. Yang, G.-Y. Zheng, GJA8 missense mutation disrupts hemichannels and induces cell apoptosis in human lens epithelial cells. Sci. Rep. 9, 19157 (2019).

42. P. J. Minogue, J.-J. Tong, A. Arora, I. Russell-Eggitt, D. M. Hunt, A. T. Moore, L. Ebihara, E. C. Beyer, V. M. Berthoud, A Mutant Connexin50 with Enhanced Hemichannel Function Leads to Cell Death. Invest. Ophthalmol. Vis. Sci. 50, 5837–5845 (2009).

43. A. Arora, P. J. Minogue, X. Liu, M. A. Reddy, J. R. Ainsworth, S. S. Bhattacharya, A. R. Webster, D. M. Hunt, L. Ebihara, A. T. Moore, E. C. Beyer, V. M. Berthoud, A novel GJA8 mutation is associated with autosomal dominant lamellar pulverulent cataract: further evidence for gap junction dysfunction in human cataract. J. Med. Genet. 43, e2–e2 (2006).

44. V. Berry, A. Ionides, N. Pontikos, I. Moghul, A. T. Moore, R. A. Quinlan, M. Michaelides, Whole Exome Sequencing Reveals Novel and Recurrent Disease-Causing Variants in Lens Specific Gap Junctional Protein Encoding Genes Causing Congenital Cataract. Genes 11, 512 (2020).

45. X.-L. Ge, Y. Zhang, Y. Wu, J. Lv, W. Zhang, Z.-B. Jin, J. Qu, F. Gu, Identification of a Novel GJA8 (Cx50) Point Mutation Causes Human Dominant Congenital Cataracts. Sci. Rep. 4, 4121 (2014).

46. A. Jin, Q. Zhao, S. Liu, Z. Jin, S. Li, M. Xiang, M. Zeng, K. Jin, Identification of a New Mutation p.P88L in Connexin 50 Associated with Dominant Congenital Cataract. Front. Cell Dev. Biol. 10 (2022).

47. J. D. Pal, V. M. Berthoud, E. C. Beyer, D. Mackay, A. Shiels, L. Ebihara, Molecular mechanism underlying a Cx50-linked congenital cataract. Am. J. Physiol.-Cell Physiol. 276, C1443–C1446 (1999).

48. A. Shiels, D. Mackay, A. Ionides, V. Berry, A. Moore, S. Bhattacharya, A missense mutation in the human connexin50 gene (GJA8) underlies autosomal dominant “zonular pulverulent” cataract, on chromosome 1q. Am. J. Hum. Genet. 62, 526–532 (1998).

49. V. Vanita, J. R. Singh, D. Singh, R. Varon, K. Sperling, A mutation in GJA8 (p.P88Q) is associated with “balloon-like” cataract with Y-sutural opacities in a family of Indian origin. Mol. Vis. 14, 1171–1175 (2008).

50. C. Sellitto, L. Li, T. W. White, Connexin50 Is Essential for Normal Postnatal Lens Cell Proliferation. Invest. Ophthalmol. Vis. Sci. 45, 3196–3202 (2004).

51. C. Sellitto, T. W. White, Combinatorial genetic manipulation of Cx50, PI3K and PTEN alters postnatal mouse lens growth and homeostasis. Front. Ophthalmol. 5 (2025).

52. T. W. White, Y. Gao, L. Li, C. Sellitto, M. Srinivas, Optimal Lens Epithelial Cell Proliferation Is Dependent on the Connexin Isoform Providing Gap Junctional Coupling. Invest. Ophthalmol. Vis. Sci. 48, 5630–5637 (2007).

53. M. Bradic, H. Teotónio, R. L. Borowsky, The Population Genomics of Repeated Evolution in the Blind Cavefish Astyanax mexicanus. Mol. Biol. Evol. 30, 2383–2400 (2013).

54. W. Sun, X. Xiao, S. Li, X. Guo, Q. Zhang, Mutational screening of six genes in Chinese patients with congenital cataract and microcornea. Mol. Vis. 17, 1508–1513 (2011).

55. J.-J. Tong, U. Khan, B. G. Haddad, P. J. Minogue, E. C. Beyer, V. M. Berthoud, S. L. Reichow, L. Ebihara, Molecular mechanisms underlying enhanced hemichannel function of a cataract-associated Cx50 mutant. Biophys. J. 120, 5644–5656 (2021).

56. R. Li, X. Wang, C. Bian, Z. Gao, Y. Zhang, W. Jiang, M. Wang, X. You, L. Cheng, X. Pan, J. Yang, Q. Shi, Whole-Genome Sequencing of Sinocyclocheilus maitianheensis Reveals Phylogenetic Evolution and Immunological Variances in Various Sinocyclocheilus Fishes. Front. Genet. 12, 736500 (2021).

57. F. Villanelo, P. J. Minogue, J. Maripillán, M. Reyna-Jeldes, J. Jensen-Flores, I. E. García, E. C. Beyer, T. Pérez-Acle, V. M. Berthoud, A. D. Martínez, Connexin channels and hemichannels are modulated differently by charge reversal at residues forming the intracellular pocket. Biol. Res. 57, 31 (2024).

58. X. Fang, I. Seim, Z. Huang, M. V. Gerashchenko, Z. Xiong, A. A. Turanov, Y. Zhu, A. V. Lobanov, D. Fan, S. H. Yim, X. Yao, S. Ma, L. Yang, S.-G. Lee, E. B. Kim, R. T. Bronson, R. Šumbera, R. Buffenstein, X. Zhou, A. Krogh, T. J. Park, G. Zhang, J. Wang, V. N. Gladyshev, Adaptations to a Subterranean Environment and Longevity Revealed by the Analysis of Mole Rat Genomes. Cell Rep. 8, 1354–1364 (2014).

59. R. Partha, B. K. Chauhan, Z. Ferreira, J. D. Robinson, K. Lathrop, K. K. Nischal, M. Chikina, N. L. Clark, Subterranean mammals show convergent regression in ocular genes and enhancers, along with adaptation to tunneling. eLife 6, e25884 (2017).

60. M. S. Springer, C. A. Emerling, J. Gatesy, Three Blind Moles: Molecular Evolutionary Insights on the Tempo and Mode of Convergent Eye Degeneration in Notoryctes typhlops (Southern Marsupial Mole) and Two Chrysochlorids (Golden Moles). Genes 14, 2018 (2023).

61. J. A. Flores, B. G. Haddad, K. A. Dolan, J. B. Myers, C. C. Yoshioka, J. Copperman, D. M. Zuckerman, S. L. Reichow, Connexin-46/50 in a dynamic lipid environment resolved by CryoEM at 1.9 Å. Nat. Commun. 11, 4331 (2020).

62. J. Jumper, R. Evans, A. Pritzel, T. Green, M. Figurnov, O. Ronneberger, K. Tunyasuvunakool, R. Bates, A. Žídek, A. Potapenko, A. Bridgland, C. Meyer, S. A. A. Kohl, A. J. Ballard, A. Cowie, B. Romera-Paredes, S. Nikolov, R. Jain, J. Adler, T. Back, S. Petersen, D. Reiman, E. Clancy, M. Zielinski, M. Steinegger, M. Pacholska, T. Berghammer, S. Bodenstein, D. Silver, O. Vinyals, A. W. Senior, K. Kavukcuoglu, P. Kohli, D. Hassabis, Highly accurate protein structure prediction with AlphaFold. Nature 596, 583–589 (2021).

63. E. F. Pettersen, T. D. Goddard, C. C. Huang, E. C. Meng, G. S. Couch, T. I. Croll, J. H. Morris, T. E. Ferrin, UCSF ChimeraX: Structure visualization for researchers, educators, and developers. Protein Sci. Publ. Protein Soc. 30, 70–82 (2021).

64. J. Liu, M. A. Riquelme, Z. Li, Y. Li, Y. Tong, Y. Quan, C. Pei, S. Gu, J. X. Jiang, Mechanosensitive collaboration between integrins and connexins allows nutrient and antioxidant transport into the lens. J. Cell Biol. 219, e202002154 (2020).

65. Y. Tong, G. Wang, M. A. Riquelme, Y. Du, Y. Quan, J. Fu, S. Gu, J. X. Jiang, Mechano-activated connexin hemichannels and glutathione transport protect lens fiber cells against oxidative insults. Redox Biol. 73, 103216 (2024).

66. B. Yue, B. G. Haddad, U. Khan, H. Chen, M. Atalla, Z. Zhang, D. M. Zuckerman, S. L. Reichow, D. Bai, Connexin 46 and connexin 50 gap junction channel properties are shaped by structural and dynamic features of their N-terminal domains. J. Physiol. 599, 3313–3335 (2021).

67. R. Jaradat, X. Li, H. Chen, P. B. Stathopulos, D. Bai, The Hydrophobic Residues in Amino Terminal Domains of Cx46 and Cx50 Are Important for Their Gap Junction Channel Ion Permeation and Gating. Int. J. Mol. Sci. 23, 11605 (2022).

68. N. C. Bennett, C. G. Faulkes, African Mole-Rats: Ecology and Eusociality (Cambridge University Press, 2000).

69. E. A. Banks, M. M. Toloue, Q. Shi, Z. J. Zhou, J. Liu, B. J. Nicholson, J. X. Jiang, Connexin mutation that causes dominant congenital cataracts inhibits gap junctions, but not hemichannels, in a dominant negative manner. J. Cell Sci. 122, 378–388 (2009).

70. Y. Shi, X. Li, J. Yang, Mutations of CX46/CX50 and Cataract Development. Front. Mol. Biosci. 9 (2022).

