## Supplemental materials for "Long-term hybridization in a karst window reveals the genetic basis of eye loss in cavefish"

### Materials and Methods

Study system

The entrance of Caballo Moro (CM) cave arose when the roof of an enclosed cave collapsed (Fig. S1). The resulting karst window allows sun exposure to a portion of an 18 x 90 m pool in the cave, which contains *A. mexicanus* cavefish. The CM cave has no known surface water basins that drain into the cave, nor any nearby resurgences (*1*). The closest surface *A. mexicanus* population is 4 km away (*2*). Loose limestone rocks and boulders from the roof collapse suggests that the karst window is a geologically recent event (Fig. S1; *3*).

Permits

Permits for wild fish collections from Secretaría del Medio Ambiente y Recursos Naturales, México, were issued to Patricia Ornelas García.

Original Caballo Moro collections: SGPA/DGVS/2438/15-16

Subsequent Caballo Moro collections: SGPA/DGVS/05389/17-18

Calera collections: SPARN/DGVS/06514/23

Animal care

All animal housing, husbandry, and experiments were conducted pursuant to Institutional Animal Care guidelines under the supervision of the Stowers Institute for Medical Research protocols 2022-129 and 2022-139.

Animal collection

Caballo Moro cavefish were collected in May 2016 and January 2017, including 17 light-dwelling eyed fish, 19 dark-dwelling eyeless fish. Three surface fish each were collected from Coahuila and La Servilleta rivers (22°52’11”N 99°14’46”W; 22°50'54"N 99°7'23”W). All fish were assayed for aggression, imaged, fin clipped, and released at their original location. Five eyeless and six eyed Caballo Moro fish collected from a previous study (*2*) were used for analysis of skull formation.

For correlation of *cx50* variants with eye size and body condition, 121 specimens from the Calera cave were captured with hand nets in Dec 2022-Jan 2023 and Nov 2023 and were released immediately after measurements. Images of both eyes per fish were scored on a scale of 1-5 (see the section on Cx50 in *A. mexicanus* below for phenotyping criteria). To obtain DNA samples, we used a less invasive procedure, modified from Legendre et al., (2023). All individuals were rapidly released in their natural pools after sampling.

Skull morphology

In some cavefish individuals, the suborbital bones IV and V may be fused, the supraorbital bone is enlarged, and there is a substantial gap between the surrounding bones (preopercular and opercular) compared to surface fish (*5*). Both sides of the face were analyzed since bone asymmetries are common. Fish were cleared and double stained for cartilage and bone by the Alcian Blue-Alizarin Red method (*6*).

Aggression assay

Observations of the Caballo Moro cave were performed to corroborate the original observation of eyed fish nipping and chasing blind fish on the illuminated side by Espinasa & Borowsky (2000). We conducted a resident-intruder assay in the field, similar to Elipot et al., (2013) (Fig. S2). A fish tank was partitioned with two glasses so that there were three compartments; two small compartments 5 cm wide on the outer borders of the tank and a central compartment, 40 cm wide. In the central compartment an eyed CM cavefish is deposited ("resident" fish). After 5 min acclimatization, two "intruder" fish were added. One is a less pigmented, eyeless CM cavefish on one side and on the other a pigmented, eyed surface fish. Number of strikes on the glass directed towards each intruder fish are counted during a period of five minutes. In a second trial, the intruder fish were switched to opposite sides and the number of resident fish strikes were averaged between the two trials. A total of six eyed CM fish were tested. The same test was performed for nine surface fish acting as the resident fish, collected at the Rascón locality in the field. Total number of attacks was tested with the binomial test against the likelihood that the resident fish attacks depigmented or pigmented intruders equally. The relative number of strikes between groups was compared with a Fisher's exact test.

Sequencing

DNA was extracted from Caballo Moro and surface fin clips with a Qiagen Blood and Tissue kit. Whole genome sequencing was conducted in three batches matching three individual collections. In each sequencing event, both eyed and eyeless Caballo Moro fish were sequenced such that no batch effect is expected to impact allele frequency among groups. For the first two sequencing events, which included an even distribution of eyeless and eyed samples, and all surface samples, library prep was done with Sciclone KAPA HTP 48-barcode Kit Bioo Scientific kit and were sequenced with seven total flow cells of paired-end 100-bp Illumina HiSeq-Rapid 2500. For the third sequencing event, which included eyed and eyeless samples, library prep was done with MiSeq Reagent v2 Chemistry 500 Cycle kit and sequenced with a single flow cell for paired-end 250-bp Illumina MiSeq system. Libraries were demultiplexed with Illumina bcl2fastq2 v2.18, and quality control was carried out with an in-house pipeline including adapter-trimming (Trimmomatic v0.3091) and Fastqc. We specified a minimum quality score of 30 across a 6 bp sliding window and discarded reads with a length of <40 nucleotides. See NCBI PRJNA841601 for SRA access (Table S6).

Alignment and variant calling

Reads were aligned to the surface *Astyanax mexicanus* genome (GCA_023375975.1; *8*) using bwa-mem2 v.2.2.1. We used Picard v 3.1.1 to remove duplicates and used samtools v1.19 to split de-duplicated bams into mapped and unmapped reads. We conducted variant calling following the GATK Best Practices. Mapped bams were used to generate per-individual gvcfs with the Genome Analysis Tool Kit (GATK) v4.4.0.0 HaplotypeCaller tool (*9*). We used the GenotypeGVCFs tool in GATK v4.4.0.0 to produce vcf files for each chromosome and unplaced scaffolds that include all individuals (and include invariant sites). The SelectVariants and VariantFiltration tools in GATK v4.4.0.0 were used to apply hard filters. We then used the CombineGVCFs tool in GATK v4.4.0.0 to combine all subset VCFs for each chromosome and unplaced scaffold. This resulted in retaining a total of 1,305,420,787 sites throughout the genome,

36,053,494 of which were biallelic sites (SNPs and indels) and 25,620,339 (1.96%) of those were SNPs. All scripts used in QC, genotyping, and analysis are available at https://github.com/rikellermeyer/Caballo_Moro.

Population structure and ancestry analysis

*PCA.* For principal component analysis, biallelic sites were pruned for linkage equilibrium (r2 < 0.1) using a sliding window of 50 SNP windows, shifting by 10 SNP per step using PLINK v1.9.0 (*10*). We excluded SNPs on unplaced scaffolds and filtered to sites with a maximum of 2 no-calls in all individuals using PLINK v1.9.0. We used PLINK to prune biallelic sites to 50bp windows. Centroids were calculated with k-means clustering of PC1 and PC2 using 3 clusters, and Euclidean distance was measured between centroids with a custom R script (Fig. S3 https://github.com/rikellermeyer/Caballo_Moro/tree/main/reports/CM_graphics.Rmd). For unbiased clustering of PCA results, we used the R package factoextra to determine the optimal number of clusters and plot each cluster. For an expanded analysis, we included six genomes from Molino cavefish for admixture analysis.

*Phylogeny*. Simplified (Fig. 1) and expanded (Fig. S4) phylogeny was subset from Moran et al., (2023), which inferred a multi-species-coalescent tree from 246 individuals across 27 *Astyanax* populations.

*Population summary statistics*. We calculated summary statistics, including absolute genetic divergence (Dxy) between each pair of populations (surface, eyed, and eyeless) using non-overlapping 50kb windows with the python script popgenWindows.py (Table S1; https://github.com/simonhmartin/genomics_general/blob/master/popgenWindows.py).

*Admixture analysis*. We included a reference genome from a Molino cavefish for admixture analysis (GenBank: GCA_023375845.1). We used the dataset created by the plink filtering from the PCA analysis to conduct unsupervised ADMIXTURE analysis (*12*). ADMIXTURE v1.3.0 was run on 50, 100, 150, and 300 bp sliding windows for 1-5 ancestral populations. Cross-validation was run for K 1-5 where k represents the number of ancestral populations. Because we have a general hypothesis that there are two to three ancestral populations, the program was run on K (1-5) to widely cover the range of biologically reasonable ancestral populations.

Demographic Modeling

*Site frequency spectra*. To produce a filtered VCF containing only suitable variants for demographic modeling, we retained only SNP variants and invariant sites and excluded sites within a ±3bp buffer of an indel and within repetitive regions of the genome. Additionally, we removed sites that were heterozygous (i.e., 0/1) in every individual sampled, as these likely represent paralogous alignment from the ancient teleost genome duplication (*13*) rather than true heterozygous sequence variants. In sum, our final VCF contained only high confidence biallelic SNPs and invariant sites, as required for downstream analysis.

We generated an unfolded joint site frequency spectrum (SFS) of derived alleles for each population pair using a custom script (https://github.com/robackem/Cavefish_DemographicModeling). For all pairwise population comparisons, *Astyanax nicaraguensis* was used as the estimated ancestral state (NCBI accessions SRR9897633; SRR9897634). Sites were excluded from the SFS if they were heterozygous (i.e., 0/1) in the outgroup or if there was not a consensus between *Astyanax nicaraguensis* individuals, given that we could not determine which allele is the ancestral state at these sites. Additionally, we excluded sites with any missing genotype information in either of the *Astyanax mexicanus* populations being analyzed (i.e., for a site to have been included it must have had a genotype call in all individuals across both populations).

*Fastsimcoal2.* We performed demographic modelling using fastsimcoal2 v2.8.0.0 (*14*). Under each demographic model, the multinomial likelihood is estimated by approximating the observed SFS through coalescent simulations given a set of parameter values. The maximum likelihood estimate is achieved through an expectation conditional maximization (ECM) algorithm that iteratively finds parameters maximizing the likelihood of the given model. For all model parameters, the initial value was randomly sampled from a wide search range of potential values. In addition to estimated parameters, all models use a mutation rate of $5.62 \times 10^{-9}$ bp per generation, as measured in *Cyprinus carpio*, a closely related fish species (*15*).

For each model, we performed 100 independent fastsimcoal2 runs. We optimized parameter estimates through 100 ECM cycles and the SFS was estimated through 250,000 coalescent simulations. The rigor of our parameter estimation meets or exceeds that of established studies (e.g., Marchi et al., 2024; Rosser et al., 2024). After 100 runs of a model had completed, we calculated Akaike information criterion (AIC) values from the maximum observed likelihood achieved in each run. After AIC was calculated for all runs of all models, we converted the AIC values into an Akaike weight for each overall model. The Akaike weight serves as a value that could be directly interpreted as the conditional probability for each model (*18*). In other words, the Akaike weight reflects the probability that the given demographic model best replicates the observed SFS among the models tested. Subsequently, we selected the models with the highest Akaike weight as the best-fitting models. All demographic models and related scripts for running fastsimcoal2 can be found at https://github.com/robackem/Cavefish_DemographicModeling.

We completed a Shapiro-Wilk test for normality on the parameter estimates from our best models and determined that the data are normally distributed ($p > 0.05$). Therefore, we summarized the predicted parameter values as mean value taken from the ten runs with the best maximum estimated likelihood for the given model.

Candidate gene selection

*Genome-wide association studies.* We used GWAS to assay for loci associated with eye loss. From the VCF file generated with PCA filtering above, we used PLINK v1.9.0 to conduct a Cochran-Armitage trend test between the eyeless subpopulation and eyed fish (both eyed Caballo Moro fish and nearby surface fish). Biallelic sites were filtered to a significance of $-\log10(p)>9$, and included 10,091 variants and 467 genes, were annotated with the GTF file for AstMex3_surface from NCBI (Table S2).

*Genotype filtering and variant annotation.* We sought genetic variants that contribute to the phenotypes specific to eyeless cavefish and not to eyed cavefish or surface fish. We filtered VCF files to SNPs with 0 nocalls across all individuals and split the VCF into eyed, eyeless, and surface individuals with SelectVariants from GATK v4.4.0.0. With SelectVariants, we filtered for alleles in eyeless fish that are homozygous for the alternate allele (A/A) compared to the surface reference (R/R). Eyed fish were filtered to either heterozygous or homozygous for the reference allele (R/A or R/R). We used BCFtools v1.19 isec command to intersect alleles that match these criteria. This resulted in 203 SNPs. SNPs were annotated with SnpEff v5.2 (*19*) and the GUI interface to Ensembl's VEP (version 103). Only 87 of the 203 SNPs matched to annotated genes, spanning 40 genes (Table S3).

*Candidate gene selection*. To link eyeless Caballo Moro alleles to eye-specific genes, we developed a database of potential eye phenotype genes from previously identified QTL regions (*20*) and a custom *D. rerio* gene ontology (GO) term database. Wiese et al., (2025) compiled 12 studies that evaluated eye phenotypes for QTLs in cave-surface F2 hybrids and identified 58 QTL genomic regions that associate with the following phenotypes: eye area, eye diameter, eye orbit area, eye size, lens, ossified sclera, pupil diameter, and pupil lens area. Some of the 58 QTLs are overlap. We used a custom python script to match genes from the AstMex3_surface GTF file to eye QTL regions, which encompasses 15,233 potential genes (Table S4; https://github.com/rikellermeyer/Caballo_Moro/candidate_gene_selection/qtl_to_genes.py). We generated a zebrafish database of genes with eye associated gene ontology terms using AmiGo v2.5.17 and PANTHER v19.0. To construct this database, we searched for zebrafish genes with gene ontology terms that associate with the biological processes: eye, lens, optic, retina, sclera, vision, and visual. The zebrafish GO term database included 810 eye-associated genes (Table S4).

Finally, we compiled a list of genes under a hard selective sweep in at least one cave out of seven populations from Moran et al., (2023). From Moran et al., (2023) there are 12,095 genes under hard selective sweeps, which represents a region under selection in cavefish.

We narrowed down our list of candidate genes for eye loss as the intersection of: 1) homozygous for the alternate allele in eyeless fish, but not eyed or surface (genotype filtering), 2) statistically associates with eye loss in the eyeless Caballo Moro fish (GWAS), 3) has undergone a hard selective sweep in at least one cave population, 4) falls within an eye phenotype QTL, and 5) has known eye-related homologs in zebrafish (Fig. S7). We used a custom python script to intersect the genes identified from each analysis - GWAS, genotype filtering, QTL, and GO term analysis (https://github.com/rikellermeyer/Caballo_Moro/candidate_gene_selection/intersect_candidate_genes.py). Five genes matched these criteria (Fig. S7).

<u>Bulk RNA-seq of the lens</u>

*Lens dissection and recovery*. The procedure used for recovering the lens was modified from operations designed previously for lens deletion and transplantation (*21*, *22*). At the stage 26-28 hours postfertilization (hpf), surface fish larvae were washed for 5 min in zebrafish ringer consisting of 116 mM NaCl (#E529-500ML: VWR international, Radnor, PA, USA), 2.9 mM KCl (#418205000; Acros organics, Geel, Belgium), 1.77 mM of CaCl 2 (#0556-500G; VWR international) and 10 mM HEPES pH 7.2 (#J848-500ML; VWR international), followed by incubating with zebrafish ringer with high calcium (116 mM NaCl, 2.9 mM KCl, 10mM CaCl 2 and 10 mM HEPES pH 7.2) for 10 min, then, incubated with calcium-free zebrafish ringer (CFZFR: 116 mM NaCl, 2.9 mM KCl, and 10 mM HEPES pH 7.2), rinsed in CFZFR (40˚C) containing 0.2% EDTA (#324503, MilliporeSigma, Burlington, MA, USA), and embedded in 1.2% agar (#BP1423-500, Fisher Scientific) in CFZFR (40˚C) by pouring the embryo containing agar into 10 cm sterile dish (#FB0875712, Fisher Scientific). After cooling to room temperature, the operations were done with sharp tungsten needles on larvae embedded in the agar.

For lens recovery, larvae were positioned on their lateral side, and small cuts were made around the lens to detach it from the ciliary body using a tungsten needle. The lens was then removed. The RNase-free BSA (#B9000; New England Biolabs, Ipswich, MA, USA) treated P10 filtered pipette tip (#02-707-456; Fisher Scientific, Waltham, MA, USA) is used to recover each lens into a 1.5 mL RNase-free microcentrifuge tube (#02-682-002, Fisher Scientific). In every lens recovery, the lenses were moved to the clean agar area (not the dissection site) and we carefully ensured the recovery was successful under the dissection microscope (SZ51 Stereo microscope with the LED oblique illumination; Olympus/Evident, Tokyo, Japan). Each tube was incubated on ice until the operation was done (6-10 fish; 45 min to 60 min). After the operation, all 1.5 mL tubes were submerged in liquid nitrogen and stored in -80˚C until shipment.

*RNA isolation and sequencing.* Once all samples were gathered and frozen, total RNA was extracted using the mirVana RNA isolation kit (Catalog number: AM1560) according to the manufacturer's instructions. cDNA was generated from 500 pg of total RNA, whose quality and quantity were assessed using a Bioanalyzer (Agilent Technologies), with 13 PCR cycles and the SMART-Seq v4 Ultra Low Input RNA Kit for Sequencing (Takara, Cat. No. 634891). Libraries were prepared from 1 ng of cDNA using 12 PCR cycles with the Nextera XT DNA Library Preparation Kit (Illumina, Cat. No. FC-131-1096), according to the manufacturer's instructions. Indexing was performed using Nextera XT Index Kit v2 (Illumina, Cat. No. FC-131-2001). Libraries were purified with AMPure XP beads (Beckman Coulter, Cat. No. A63882) and assessed for quality and concentration using a Qubit Fluorometer (Life Technologies) and Bioanalyzer. Equal molar libraries were pooled, quantified, and sequenced on a High-Output flow cell of an Illumina NextSeq 500 instrument using NextSeq Control Software v4.0.1, with the following read configuration: 76 bp Read 1 and 6 bp i7 Index. Following sequencing, Illumina Primary Analysis (NextSeq RTA v2.11.3.0) and bcl-convert v3.10.5 were used to demultiplex reads and generate FASTQ files.

FASTQ files were aligned to Astyanax_mexicanus-2.0 (GCF_000372685.2) using STAR (*23*) version 2.7.3a for read count summarization with quantMode GeneCounts and gene models from Ensembl 102. Transcripts per Million (TPM) values were calculated using RSEM (*24*).

CRISPant knockdowns

*CRISPant generation.* Functional analysis of *cx50* was performed using the CRISPR-Cas system to knockdown *cx50* at the one-cell stage. Three guide RNAs (sgRNA) were selected using CHOPCHOP v3 (*25*) and contained the following target sequences (PAM is underlined): TGTGGCAAATGTAGGTTCGGAGG, GGGCAGCGTCCGTACAGCCAAGG, and ACACCCTCACTGGTGTACGTGGG. To avoid off-target effects, including targeting of *gja8a*, we selected guides with no predicted off-target activity in the *Astyanax* genome; any potential off-target sites had three or more mismatches, which would prevent guide activity (*26*). The final RNP cocktail contained

30.5uM Alt-R™ S.p. Cas9 Nuclease V3 (IDT) and 10.17uM of each guide, purchased as Alt-R™ CRISPR-Cas9 sgRNAs (IDT). One nL of the sgRNA-Cas9 complex was injected at one cell stage in a transgenic surface fish line expressing green fluorescent protein (eGFP) ubiquitously, named SF-Ubi-GFP as described in (*27*).

*Cx50 eye size correlations*. Genotyping of *cx50* variants in *A. mexicanus* was conducted with fin clips or fry bodies. We conducted amplified the 472 base pairs surrounding the target site with PCR followed by sequencing on an Illumina MiSeq, followed by cleanup and quality control of the sequences. We identified knockouts as those with out-of-frame indels within amino acids 12 - 158, which includes transmembranes 1 and 2.

*cx50* forward primer: CTC TGA GGA AGT CAA CGA GCA CTC GA

*cx50* reverse primer: GAG TGT CTT GAA GAT GAT GTG GCA AAT GTA GG

Crispant *Cx50* knockouts and uninjected fish from the same clutch of SF-Ubi-GFP were collected at 2dpf and 6dpf fish, both eyes were imaged with a Leica M205 C, GFP filter, and Leica DFC 7000 T camera, and scored blindly on a scale of 1 – 5. A scale of eye degeneration phenotypes was used to capture the breadth of degeneration, allow for non-lethal sampling, and avoid confounding variables such as age and environmental conditions in wild caught samples (*28*). As such, we have used this 1-5 scale for all eye phenotyping. Examples of eye scoring can be found in Fig. 2e, f and Fig. S9 based on the following criteria:

Score of 5: Indistinguishable from age-matched surface fish.

Score of 4: Smaller eye diameter than surface fish or abnormal eye shape, and normal lens.

Score of 3: Intermediate sized eye outline, small or absent lens.

Score of 2: Small outline of the eye visible, no lens.

Score of 1: No lens or eye visible. Complete collapse of pigmented cells into a single source.

Cyclops phenotype was defined as having a single eye and lens or a single, fused iris (Fig. S9). Adult wild-caught fish from the Calera cave, described above, and surface x Tinaja F2 hybrids (496dpf) were scored from 1-5 from the scale depicted in Fig. 2e, f. Quantification of eye sizes was evaluated with Kruskal-Wallis test with Dunn's multiple comparisons.

*TUNEL assay*. CRISPant lens apoptosis was evaluated similar to (*29*). Briefly, injected and uninjected CRISPants were fixed in 4% PFA overnight. Whole fish were labeled with the Roche In Situ Cell Death Detection Kit, AP (catalogue # 11684809910) and Roche BCIP®/NBT solution, premixed (catalogue # 11681451001) for signal conversion. Lenses were removed for imaging.

<u>Cx50/CX50 in other organisms</u>

We used NCBI BLAST+ v2.15.0 to identify Cx50 variants from 19 non-*Astyanax* cavefish, 5 subterranean mammals, and their closest surface relatives using the zebrafish and mouse Cx50 sequences (Fig. S11, S12; Table S5). *Sinocyclocheilus sp.* (Chinese cavefish) had an additional whole-genome duplication compared to *Astyanax*, for which we evaluated both *cx50* paralogs and displayed the closest in sequence similarity. Amino acid sequences were evaluated for subterranean-specific variants in the first 245 N-terminal amino acids. Nonsynonymous mutations, specifically in regions with cataract-causing SNPs in humans, were prioritized for static protein modeling.

Protein modeling

We computationally made static models of selected Cx50/CX50 variants to evaluate the potential effect of variants on protein structure. We locally installed the colabfold (https://github.com/YoshitakaMo/localcolabfold; *30*) implementation of AlphaFold2 (*31*) to predict the hexameric structures of the N-terminal 245 residues of Connexin 50 from different species. The program was run on an NVIDIA A100 GPU. That fragment corresponds to the well predicted domain and experimentally resolved structure of Connexin 50 (Sheep PDB: 7JJP; *32*). A single prediction was made for each sequence and relaxed within the colabfold program. All predictions corresponded roughly to the expected hexameric gap junction structure. Visualizations of models were performed with ChimeraX v1.8 (*33*).

Firstly, a 140 x 140 angstrom squared membrane of 1-palmitoyl-2-oleoyl-sn-glycero-3-phosphocholine (POPC) was generated with VMD v1.9.4 (*34*) using the membrane plugin. Next waters were stripped from the membrane which was then overlayed onto the oriented *Astyanax* surface protein. Lipid molecules less than 1 angstrom from protein atoms were removed at this stage and the resulting membrane was exported as a GROMACS v2024.3 gro file (separate from the protein file; *35*). For other proteins alignment was performed to the surface protein and the membrane structure was used as is. Membrane and protein files were then converted to Gromacs format with appropriate topologies by the pdb2gmx utility with structural outputs as pdb files. Lipid topologies were converted to independent files and referenced in the protein topology file to create a combined topology file. A combined protein and membrane pdb file was created by manually concatenating protein and membrane files and removing the membrane pdb header lines. The combined file was placed into a 140 angstrom dimension cubed box using the editconf utility and solvated using the solvate utility and tip3p water. Water molecules in the membrane were removed using a custom python script, and the topology file was updated in a corresponding fashion. The solvated filtered output then had sodium and/or chloride ions added to it (with the genion utility) to achieve neutral charge.

This final preparation was submitted to energy minimization with potential energy minimizing to values approaching $-3 \times 10^6$ kJ/mol. Next a 100 ps NVT simulation was run with temperature at 323K (well above the transition point for POPC and maintained for all simulations) followed by a 1 ns NPT simulation with a Nose-Hoover thermostat. Potential, temperature, and pressure stability were verified at each step. Next 1 ns of unconstrained molecular dynamics was performed and stability was verified by visual inspection before running 5 ns of production molecular dynamics. This final 5 ns data set was used for further analysis.

Trajectory import and export was done with the MDAnalysis (https://github.com/MDAnalysis, *36*) python package and periodic boundary condition unwrapping was performed with VMD (see above). Protein and lipid analysis and visualizations were performed at 5 ps intervals while water movement was analyzed at the full output time resolution (0.2 ps). Water molecules were first filtered for those that appeared within a 15 angstrom lateral radius of the pore and 5 angstroms from the axial pore center. For water pore crossing calculations, inner, middle, and outer pore regions were defined as cylinders with lateral radius 15 angstroms and 30 angstrom height with the middle pore region centered on the channel. Crossings were counted when molecules transited from inner to middle to outer regions (outer crossings) or in the reverse order (inner crossings).

*Pore calculations.* Pore measurements were performed in python using the jpdbtools2 package well as a pore_utils package found here: https://github.com/jayunruh/Jay_pdbtools/. The *Astyanax mexicanus* surface fish predicted structure was first aligned vertically by visual inspection (as in the molecular dynamics preparation step) and used as the reference structure. Then all other well-predicted structures were aligned to that structure by least squares fitting of alpha carbon atoms. In cases where the sequences were shorter than the reference structure the alignment was only performed on the N-terminal overlapped region. Surfaces were created by projecting the molecular coordinates onto a 0.25 angstrom resolution grid and masking all grid voxels within 2.6 angstroms of the nearest atom. This corresponds approximately to the solvent accessible distance for most atoms. The center of each pore was found as the center of mass of the sum projected pore voxels in the direction of the channel masked with a 110 voxel radius cylinder. Finally, the pore radius was measured as the minimum distance from that center of mass at each voxel layer traveling through the pore.

**Mouse CX50-S89K**

*Mouse Cx50 CRISPR*. CRISPR-Cas technology was used for engineering C57BL/6J mice to carry the exact mutation found in cavefish. Potential guideRNA target sites were designed using CCTOP v1.0.0 (*37*). The target site was selected by evaluating the predicted on-target efficiency score and the off-target potential (*25*) in addition to the proximity of the double strand break to the desired mutation site. To generate the S89K codon change, a single stranded DNA oligonucleotide (ssODN) donor was designed containing ~50 nucleotides of homology from the double strand break site. The ssODN contained mutations for the S89K change as well as silent codon changes to disrupt guideRNA binding following the repair event. The ssODN was ordered as an Ultramer from Integrated DNA Technologies (IDT). The selected guideRNA was ordered as an Alt-R CRISPR-Cas9 crRNA from IDT. The crRNA was hybridized with tracrRNA at 100uM. RNP was formed using cr/tracrRNA at 6uM final concentration with IDT Cas9 HiFi v3 protein at 1.2uM final concentration in 100ul. The RNP was formed by incubating at room temperature for 10 minutes. The ssODN was added at 400ng final concentration to the RNP and delivered to LASF for electroporation into B6 mouse embryos (guide: TCGCTGATGTACGTGGGGCA).

Mice were screened for the expected mutations by lysing an ear clip and amplifying the specific genomic location. A second round of amplification was done to incorporate sample-specific dual barcodes. All amplicons were pooled and size-selected using ProNex Size-Selective Purification System (Promega). Cleaned pools were quantified on a Qubit Fluorometer and then ran on an Agilent Bioanalyzer to check sizing and purity. Purified pools were run on an Illumina MiSeq 2x250 flow cell. The resulting sequence data was demultiplexed and read pairs were joined. On-target indel frequency and expected mutations were analyzed using CRIS.py (*38*).

Mouse primers:

*Cx50* Forward primer: CTACGATGAGGCCTTTCCCA

*Cx50* Reverse primer: CTCCATGCGAACGTGGTGT

*Mouse phenotyping*. Eye samples were prepared and stained with hematoxylin & eosin (H&E; ST Infinity H&E Staining System, VWR Cat#10015-090) according to Pang et al., 2021. Diameter of eyes and lenses were taken on multiple sections and measured with ImageJ v1.46r. Comparisons were made between heterozygous, homozygous, and wild-type mice from ages 162 to 226 days only. Eye and lens size were averaged across three to five sections per eye with a least squares multiple linear regression model.

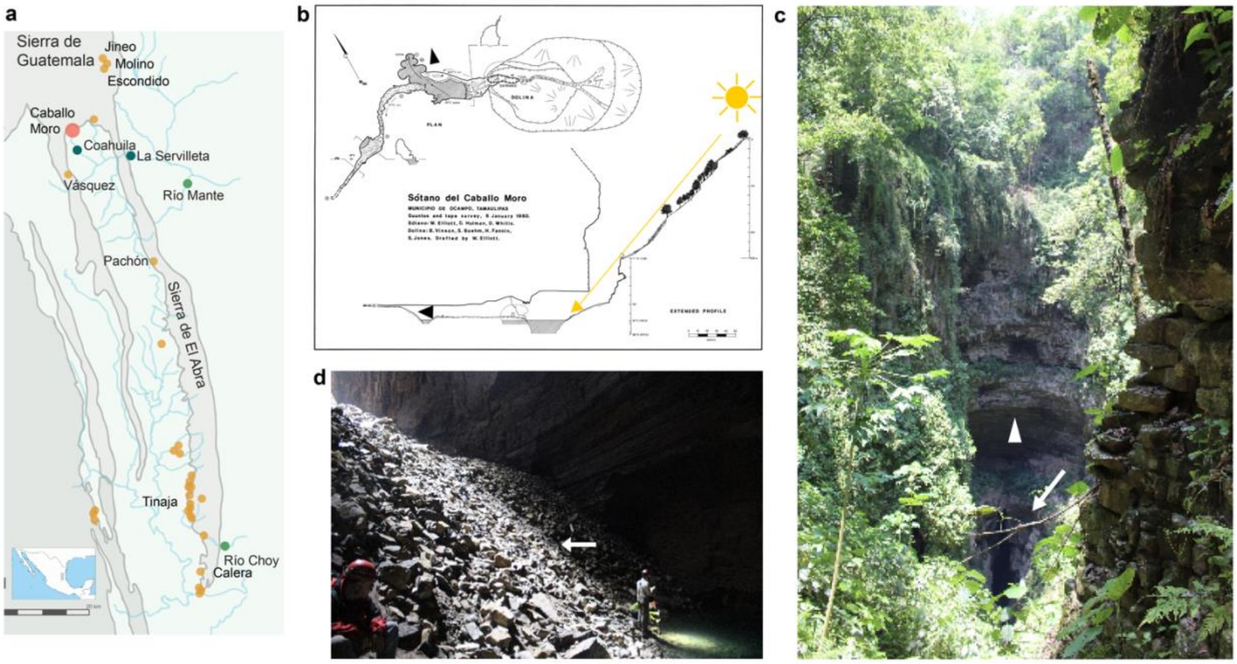

**Fig. S1. The Caballo Moro cave. (a)** Map of the Sierra de Guatemala and Sierra de El Abra regions in northeastern Mexico. Select surface fish populations are shown in green, Caballo Moro in orange (large dot), and other caves in orange (small dots). **(b)** Map of the Caballo Moro cave adapted from Elliott (2018). The lake extends further into the cave (arrowheads) and only half of the lake is exposed to sunlight. **(c)** The entrance of the Caballo Moro cave (arrow) showing the remaining roof of the karst window (arrowhead). **(d)** The base of the cave entrance has large, unweathered rocks (arrow) from the ceiling collapse with minimal evidence of wear, indicative of a relatively recent ceiling collapse.

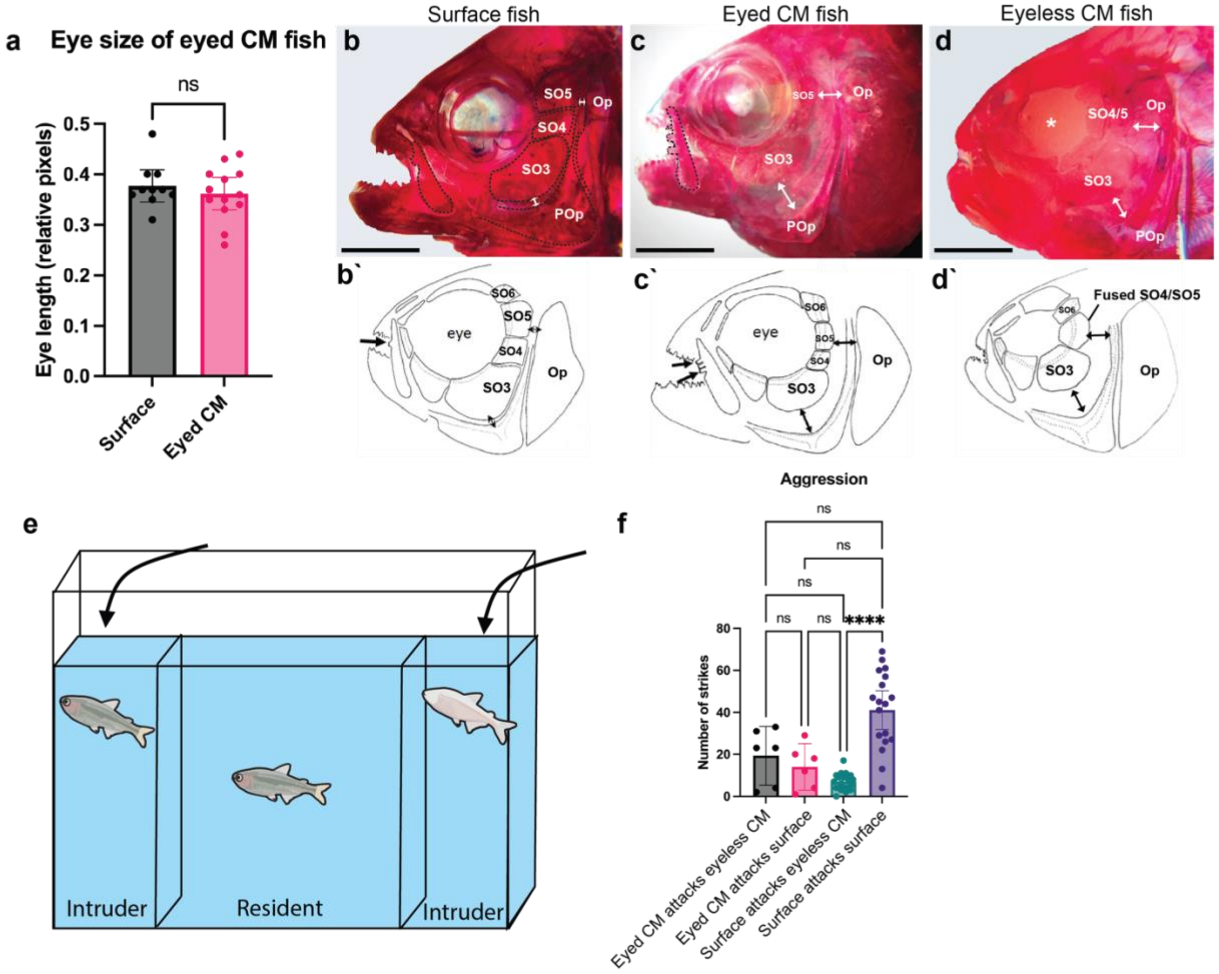

**Fig. S2. Cave-like (troglomorphic) characteristics of the eyed Caballo Moro fish and aggression. (a)** Eye diameter of Caballo Moro (CM) eyed fish (n=7) and surface fish (n=5) measured in relative pixels (both eyes measured for each individual; p=0.53, Mann-Whitney U test, error bars are 95% confidence interval). **(b-d')** Suborbital bone structure of surface and Caballo Moro fish visualized with alizarin red. **(b)** Surface fish with suborbital bones (SO3-SO5). There are small gaps between the suborbital bones and the opercle (Op) and preopercle (POp) bones (double sided arrows). Single maxillary teeth (outlined) are common in surface fish. **(c)** Eyed Caballo Moro fish have larger gaps between the SO5 and the Op and the SO3 and the POp bones. Three maxillary teeth are outlined and are typical in eyed Caballo Moro fish. **(d)** Eyeless Caballo Moro fish, have large gaps between SO5/Op and SO3/Pop and sometimes suborbital bones fused. While difficult to image, eyeless Caballo Moro fish typically have 3 maxillary teeth. **(e)** A resident-intruder assay places a “resident” fish in the center of a plexiglass arena to acclimate for 5 minutes. An intruder fish is then added to each side and the number of strikes the resident takes at each intruder were counted over 5 minutes. The intruder fish are then switched sides and counts occur for another 5 minutes. Trials were conducted with eyed Caballo Moro fish or surface fish as the resident. For both experiments, the intruders were depigmented, eyeless Caballo Moro fish and surface fish. **(f)** Quantification of the number of times the resident eyed Caballo Moro fish struck intruder fish per trial (Eyed CM attacks eyeless CM and Eyed CM attacks surface; n=6 trials). Quantification of strikes taken by surface fish against eyeless Caballo Moro fish and surface fish (n=9 trials; n=9, **** p<0.0001, two-sided Fisher's exact test, error bars are 95% confidence interval).

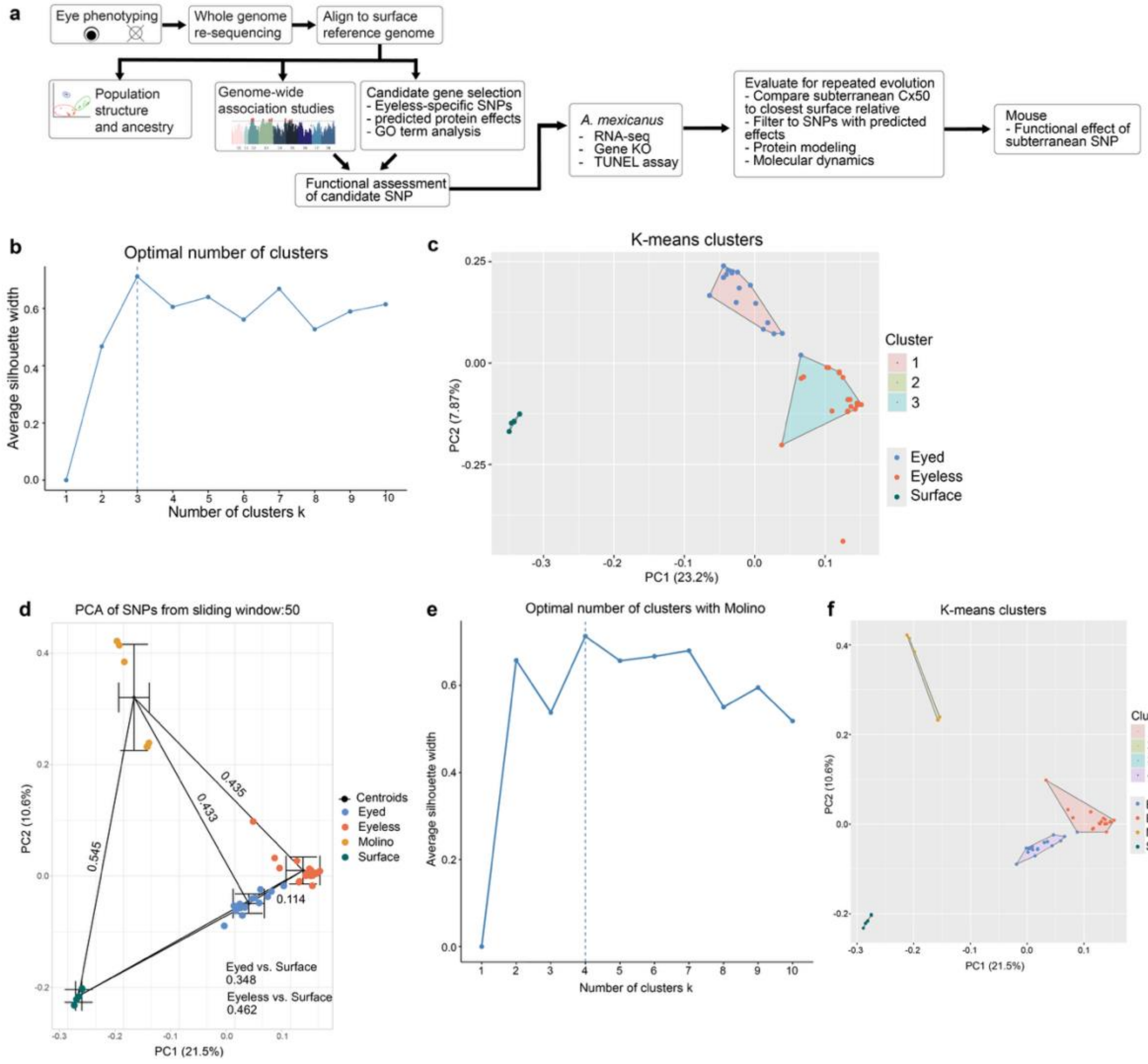

**Fig. S3**. **Caballo Moro population analysis. (a)** Workflow for Caballo Moro cavefish population and candidate gene analysis. **(b)** The optimal number of unbiased k-means clusters of eyed, eyeless, and surface fish, is indicated as the best fit of the average silhouette width (dotted line). **(c)** Unbiased k-means clustering of eyed, eyeless, and surface fish as three clusters. Note a single eyed fish clusters with eyeless and a single eyeless fish does not cluster. **(d)** Expanded PCA showing distance between mean centroids of each subpopulation (eyed, eyeless, Molino cavefish, and surface), measured as Euclidean distance. Error bars indicate standard error. **(e)** The optimal number of k-means clusters for the expanded PCA dataset that include Molino cavefish. Four clusters indicate the best fit for the data set (dotted line). **(f)** Unbiased k-means clustering of eyed, eyeless, Molino, and surface fish. Note a single eyed fish clusters with eyeless fish.

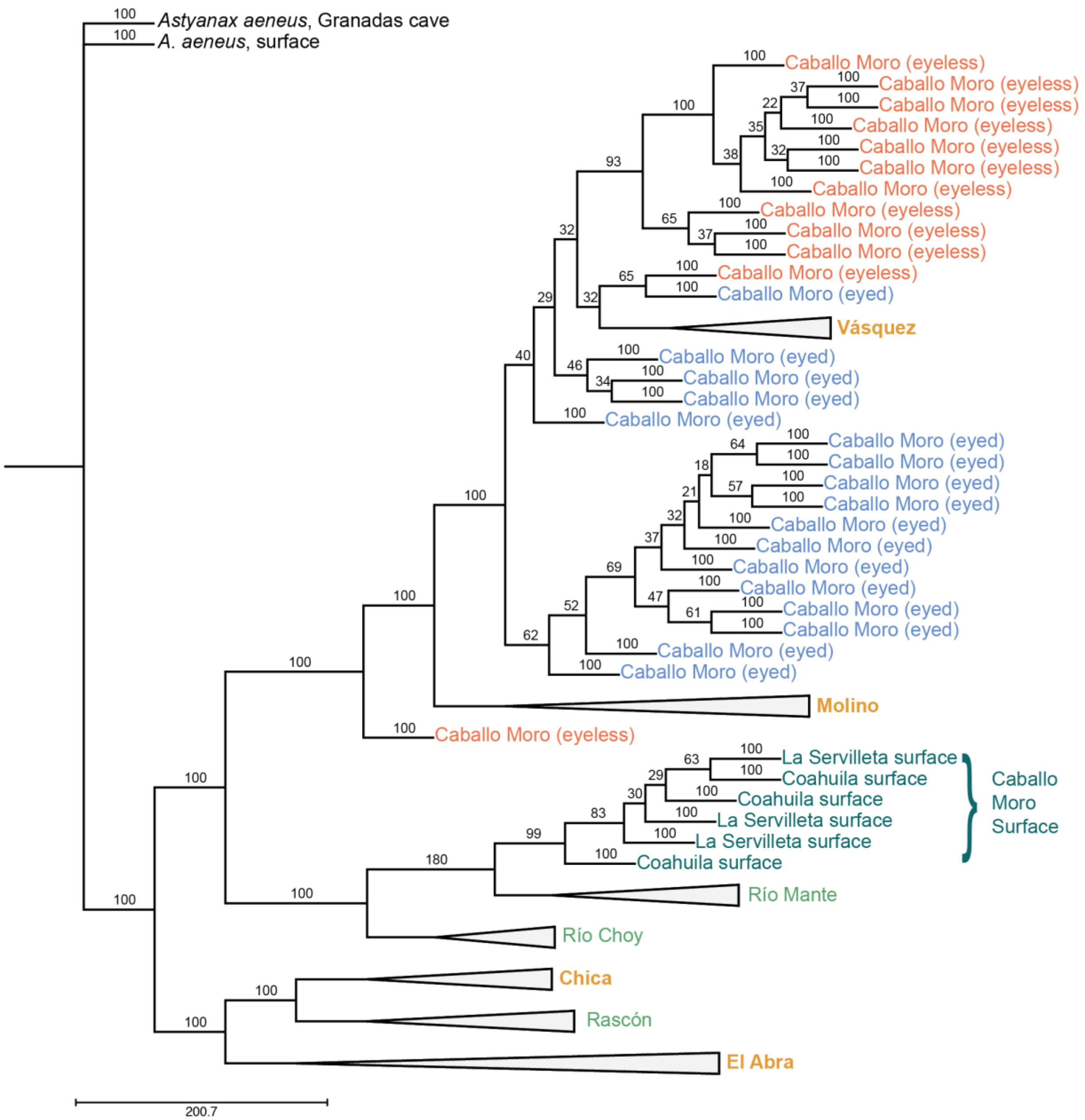

**Fig. S4. Caballo Moro phylogenetic analysis.** Expanded phylogeny of Caballo Moro cavefish, nearby surface populations, and other cave populations from the SVDQuartets tree in Moran et al. (2023).

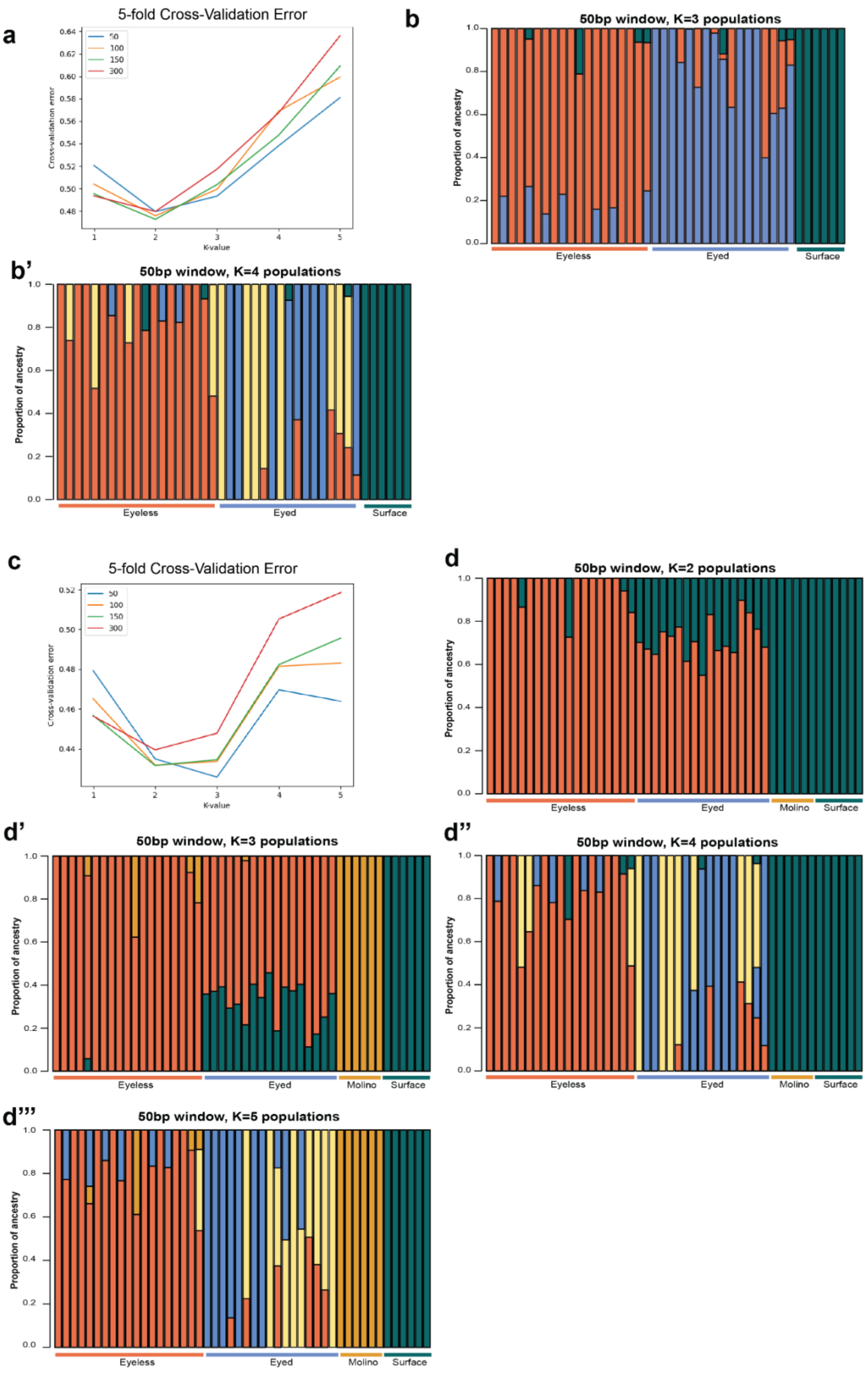
a
5-fold Cross-Validation Error
50
100
150
300
Cross-validation error
K-value
b
50bp window, K=3 populations
Proportion of ancestry
Eyeless
Eyed
Surface
b'
50bp window, K=4 populations
Proportion of ancestry
Eyeless
Eyed
Surface
c
5-fold Cross-Validation Error
50
100
150
300
Cross-validation error
K-value
d
50bp window, K=2 populations
Proportion of ancestry
Eyeless
Eyed
Molino
Surface
d'
50bp window, K=3 populations
Proportion of ancestry
Eyeless
Eyed
Molino
Surface
d''
50bp window, K=4 populations
Proportion of ancestry
Eyeless
Eyed
Molino
Surface
d'''
50bp window, K=5 populations
Proportion of ancestry
Eyeless
Eyed
Molino
Surface

**Fig. S5. Admixture analysis of Caballo Moro fish. (a-b')** Admixture analysis of eyed and eyeless Caballo Moro and surface fish. **(a)** Cross-validation error of admixture analysis with K=1-5, with 50-300 bp windows. Lines indicate the cross-validation error for each base pair window. Lowest cross-validation error indicates model with best fit. The best fit was K = 2, which was shown in Figure 1 in the main text. **(b)** Second best-fit admixture analysis is K=3, with distinct ancestry for eyed, eyeless, and surface fish. **(b')** Third best-fit admixture analysis, K=4 with 50bp windows. Surface fish and eyeless fish maintain distinct ancestries, with some evidence of recent admixture. **(c-d''')** Admixture analysis of Caballo Moro fish, surface fish, and Molino cavefish. **(c)** Cross-validation error of admixture analysis indicates that K = 3 is the best fit. **(d-d''')** Admixture plots for K = 2, 3, 4, and 5 populations. K=3 is best supported by cross-validation error and shows independent ancestry for eyeless Caballo Moro, Molino, and surface fish, with eyed Caballo Moro fish sharing eyeless and surface ancestry.

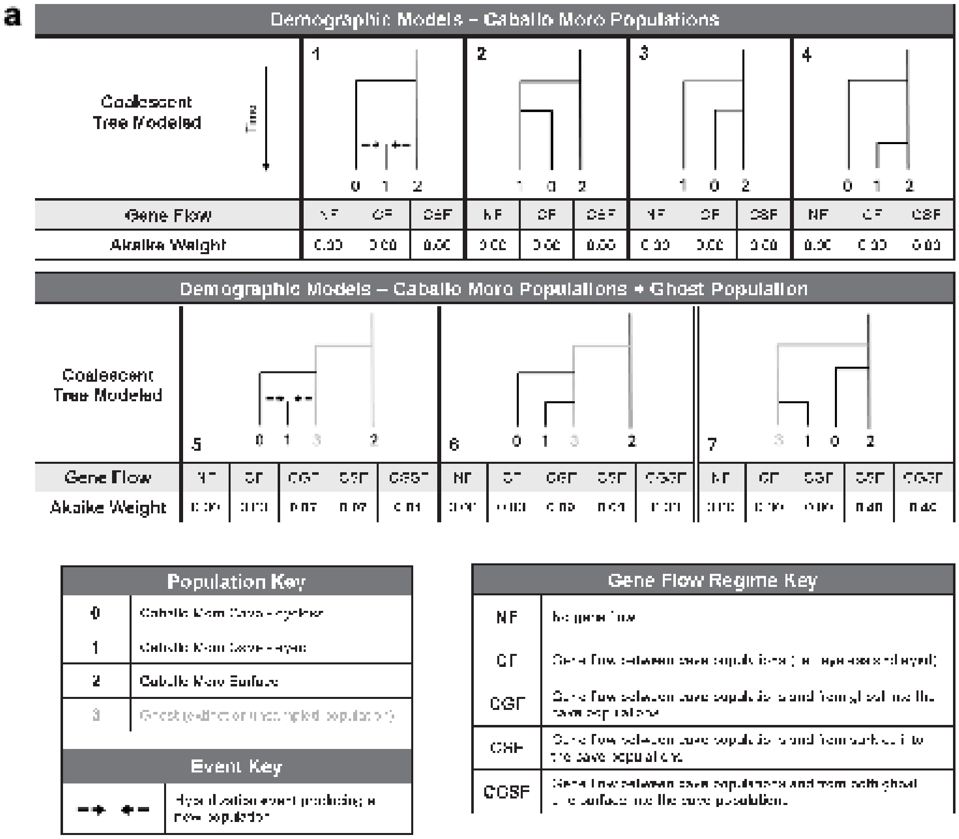

b

| Parameter Estimated | | Tree 7 CSF | Tree 7 CGSF |
|---|---|---|---|
| **Effective population size (*Ne*)** | Eyeless | 15,785 ([illegible]) | 17,582 ([illegible]) |
| | Eyed | 1,037 (29) | 1,038 (14) |
| | Surface | 522,049 ([illegible]) | 514,337 ([illegible]) |
| **Migration (*per generation*)** | Eyeless to Eyed | 1.12E-03 ([illegible]) | 1.17E-03 ([illegible]) |
| | Eyed to Eyeless | 5.08E-06 ([illegible]) | 1.39E-07 ([illegible]) |
| | Surface to Eyeless | 3.87E-05 ([illegible]) | 3.22E-05 ([illegible]) |
| | Surface to Eyed | 3.90E-04 ([illegible]) | 3.75E-04 (1.70E-05) |
| | Ghost to Eyeless | n.a | 5.41E-07 ([illegible]) |
| | Ghost to Eyed | n.a | 4.10E-06 ([illegible]) |
| **Divergence time (*# of generations*)** | Ghost / Surface | 963,611 ([illegible]) | 978,775 ([illegible]) |
| | Eyeless / Surface | 119,860 ([illegible]) | 128,622 ([illegible]) |
| | Eyed / Ghost | 3,913 ([illegible]) | 4,354 ([illegible]) |

**Fig. S6. Demographic models of Caballo Moro ancestry. (a)** Results of demographic modeling of the Caballo Moro system with Fastsimcoal2 (*14*). Akaike weights, derived from 100 runs of each model, reflect the probability that a model is the best among the set of seven models tested. For all models, the gene flow regime begins after the final divergence of populations on the tree (i.e., the divergence that has occurred most recently). Tree 7 represents the most likely demographic model that best approximated the observed site frequency spectrum, indicating that the eyed cavefish in Caballo Moro likely came from a unique surface ancestor that is either extinct or has not been sampled. **(b)** Parameters values estimated in the two best-fitting models (tree 7 topology, see (a)). Parameter values listed are the median value taken from the ten runs with the best maximum estimated likelihood for the given model, accompanied by the standard deviation in parentheses. Ne values are given as haploid number (i.e., 2*number of individuals). The migration value reflects the probability that an individual in the recipient population came from the donating population in the previous generation. Divergence time is measured in the number of generations back in time from present.

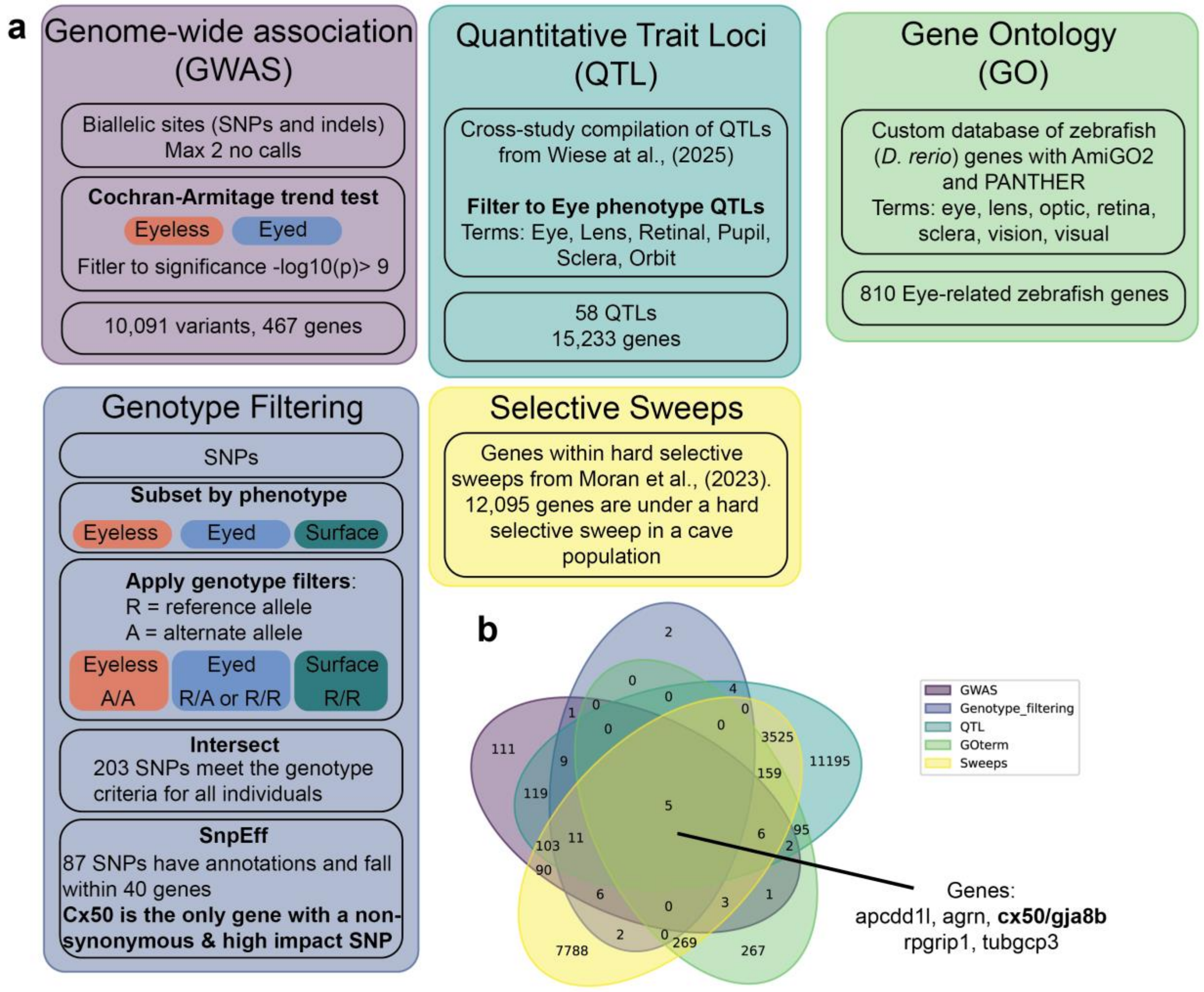

**Fig. S7. Candidate gene selection for eye loss. (a)** A graphical overview of candidate gene selection. Genotype filtering and GWAS was used to identify genes associated with the eyeless phenotype in eyeless Caballo Moro fish. Publicly available datasets - eye phenotype QTLs from Wiese et al., (2025) and zebrafish gene ontology database - were used to identify which genes associated with eyeless Caballo Moro are likely related to eye phenotypes. **(b)** Intersection of these CM datasets and eye gene databases identified five genes that meet all criteria: *apcdd1l*, *agrn*, *rpgrip1*, *tubgcp3*, and *cx50/gja8b*. Adenomatosis polyposis coli down-regulated 1-like (*apcdd1l*) is expressed in retinal epithelial cells and negatively regulates the Wnt signaling pathway during vascular remodeling and maturation in the developing retina (*41*). Agrin is a heparan sulfate proteoglycan involved in the extracellular matrix surrounding synapses in the retina. Agrin (*agrn*) interacts with *shh* and *wnt* signaling pathways to modulate optic nerve growth and eye development in zebrafish - and mutations can cause microphthalmia (*42*). Tubulin gamma complex component 3 (*tubgcp3*) is part of the tubulin complexes in centrosomes and is involved in regulating the retinal progenitor cell cycle. Removal of *tubgcp3* causes microphthalmia in zebrafish (*43*). Retinitis pigmentosa GTPase regulator interacting protein 1 (*rgrip1*) is a ciliary protein expressed in the retina, to aid in the connection between photoreceptors and cilia. Mutations in *rpgrip1* result in retinitis pigmentosa, cone-rod dystrophy, and Leber congenital amaurosis (*44*). We highlight these genes as potential candidates for future study. *Cx50*/*gja8b*, focused on in this manuscript, was the only gene with a non-synonymous mutation in a protein coding region that also has a predicted high impact effect with SnpEff.

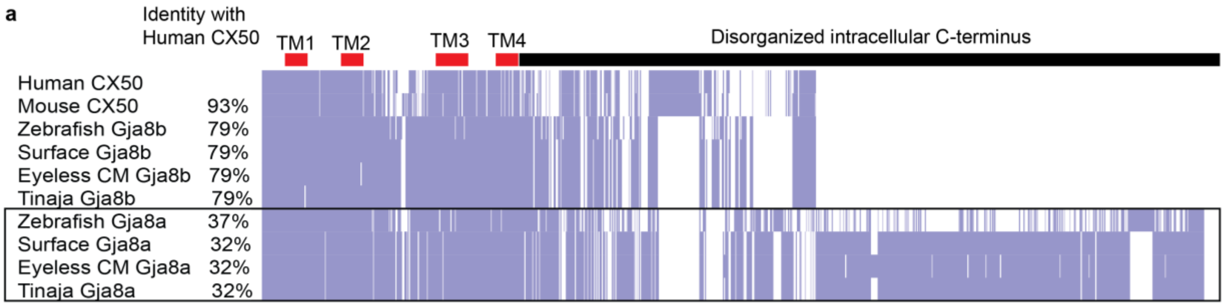

**b**

```
MGDWSFLGNILEEVNEHSTVIGRVWLTVLFIFRILILGTAAEFVWGDEQSDYVC
NTQQPGCENVCYDEAFPISHIRLWVLQIIFVSTPSLVYVGHAVHHVHMEEKRKE
REEAELSRQQEAGEERPPLAPDQGSVRTAKETSTKGSKKFRLEGTLLRTYICHI
IFKTLFEVGFVVGQYFLYGFRILPLYKCSRWPCPNTVDCFVSRPTEKTVFIIFM
LAVACVSLFLNFVEISHLGLKKIHFVFRKPARPQIEGSGSSEKPSMAVSSIQKA
KGYKLLEEDKTPSHFFPMTEVVGMEAGRLPAPYEPFVEKTNGRDSRPQEDMSKV
YDETLPSYVQTTMVGPSGACGVIRLDDDEMPVEADVDASETIEDTRPLSSLSKA
SSRARSDDLTV-
                          FVSTPSLVYVGH
       Target Seq. 1 ACACCCTCACTGGTGTACGTGGG
                          PDQGSVRTAKETST
       Target Seq. 2 GGGCAGCGTCCGTACAGCCAAGG
                          TLLRTYICHIIF
       Target Seq. 3 CCTCCGAACCTACATTTGCCACA
```

**Fig. S8. Teleost paralogs of CX50**. **(a)** BLOSUM62 sequence similarity for mammalian CX50 and teleost Gja8a and Gja8b. Both eyeless Caballo Moro sequences and Tinaja cavefish, representing the El Abra lineage, are shown. Purple bars indicate exact sequence match, light purple indicates amino acid mismatch from the consensus sequence, white space indicates gaps. Total percent identity compared to Human CX50 is on the left. Red bars indicate transmembrane domains. Black bar indicates the structurally disorganized, intracellular C-terminus. **(b)** CRISPR-Cas target sequences of Cx50 for knockout in *A. mexicanus*. Upper is the total protein sequence of Cx50, lower is the DNA sequence of guide RNAs. Red sequences indicate transmembranes. Yellow highlights indicate target sequences. Blue highlight indicates the SNP target of the S89K mutation in eyeless CM fish. Bold sequence indicates the PAM sites.

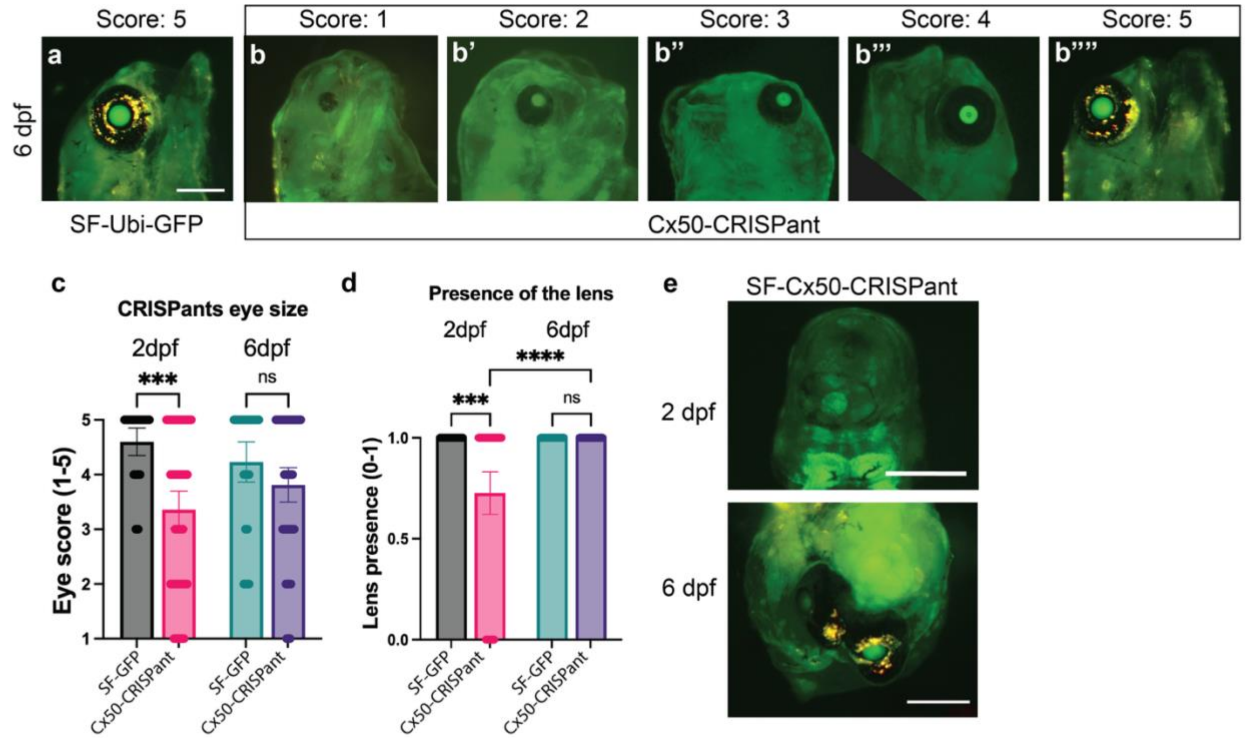

**Fig. S9. *Cx50*-CRISPants at 6dpf. (a-b'''')** Transgenic surface fish expressing Ubiquitin-GFP. Ubi-GFP allows easy visualization of the lens. Bar is 500 µm. **(a)** Normal surface eye development at 6dpf with pigment and large lens (n=20, median=5). **(b-b'''')** SF-Ubi-GFP *cx50*-CRISPants with a spectrum of eye score examples from 1-5 from 6dpf (n=35, median=4). **(c)** Comparison of *cx50*-CRISPant eye size at 2dpf and 6dpf. 6dpf eye size comparison is marginally nonsignificant p=0.085, *** indicates the p=0.0002 (Kruskal-Wallis test, error bars are 95% confidence interval). **(d)** Quantification of the presence or absence of the lens at 2dpf and 6dpf. *** p=0.0001; **** p=0.0002 (Kruskal-Wallis test, error bars are 95% confidence interval). **(e)** Transverse view of the larval head shows complete or partial cyclops phenotype with eye or eyes migrated medially. ~5% of *cx50*-CRISPants at 2dpf and 6dpf have a cyclops phenotype (n=4).

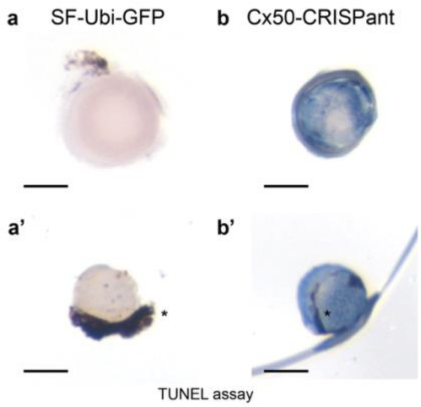

**Fig. S10. TUNEL labeling of Cx50 CRISPants surface fish lens at 6 dpf.** Lenses from 6dpf fish labeled with TUNEL (blue). Due to limited numbers of available fish, our sample size is limited and prevents quantification of TUNEL labeled cells. Note some residual melanated pupil tissue remains (asterisks). Bar is 1mm. (a) Dissected lenses of uninjected SF-Ubi-GFP fish labeled for apoptosis with TUNEL (n=6 eyes, 3 fish) **(b, b')** Dissected lenses of Cx50-CRISPants in a SF-Ubi-GFP background (n=12 eyes, 6 fish). This qualitatively demonstrates increased lens cell death in CRISPant surface fish lens.

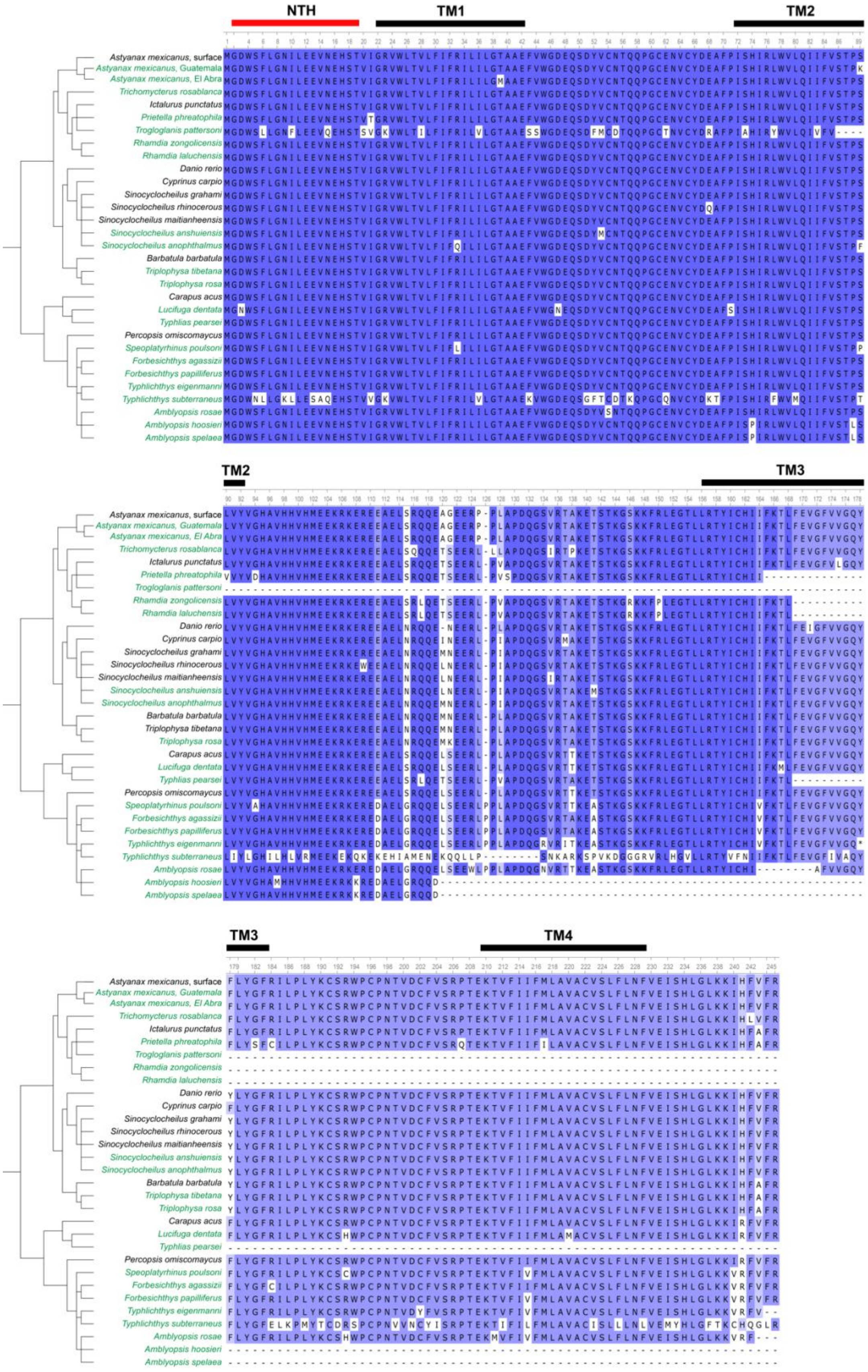
NTH
TM1
TM2
TM3
TM4
Astyanax mexicanus, surface
Astyanax mexicanus, Guatemala
Astyanax mexicanus, El Abra
Trichomycterus rosablanca
Ictalurus punctatus
Prietella phreatophila
Trogloglanis pattersoni
Rhamdia zongolicensis
Rhamdia laluchensis
Danio rerio
Cyprinus carpio
Sinocyclocheilus grahami
Sinocyclocheilus rhinocerous
Sinocyclocheilus maitianheensis
Sinocyclocheilus anshuiensis
Sinocyclocheilus anophthalmus
Barbatula barbatula
Triplophysa tibetana
Triplophysa rosa
Carapus acus
Lucifuga dentata
Typhlias pearsei
Percopsis omiscomaycus
Speoplatyrhinus poulsoni
Forbesichthys agassizii
Forbesichthys papilliferus
Typhlichthys eigenmanni
Typhlichthys subterraneus
Amblyopsis rosae
Amblyopsis hoosieri
Amblyopsis spelaea

**Fig. S11. Comparison of Cx50 sequences between cave and surface fishes.** Sequences were pulled from genome accessions available on NCBI. N-terminal helix and transmembrane domains are indicated by bars above amino acid position. Eyeless cave species are colored in green. Purple indicates level of sequence conservation. Note that not every species has a complete sequence, which influences sequence conservation calculation. See Table S5 for a table of scientific and common names.

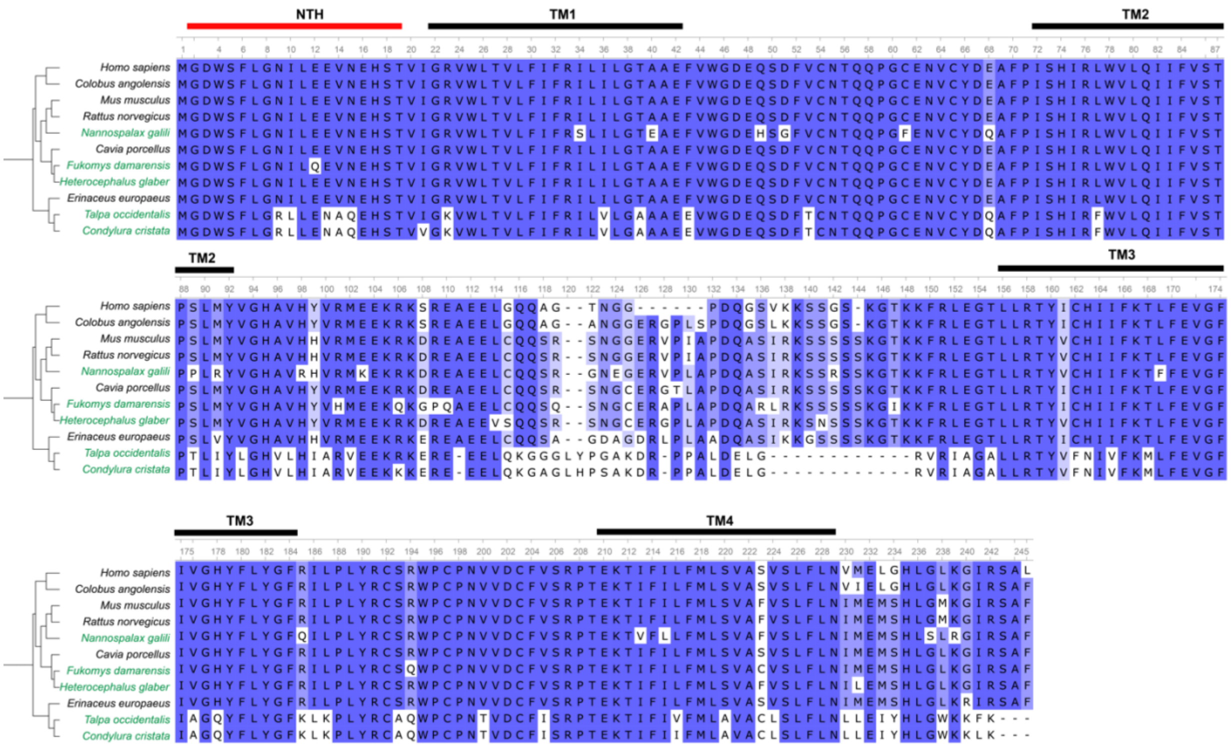

**Fig. S12. Comparison of CX50 sequences between subterranean and surface mammals.** Sequences were pulled from genome accessions available on NCBI. N-terminal helix (NTH) and transmembrane (TM) domains are indicated by bars above amino acid position. Subterranean species are colored in green. Purple indicates level of sequence conservation. Note the differences in sequence between subterranean animals and surface relatives in the NTH and transmembranes 1 and 2 as especially relevant for CX50 structure. See Table S5 scientific and common names.

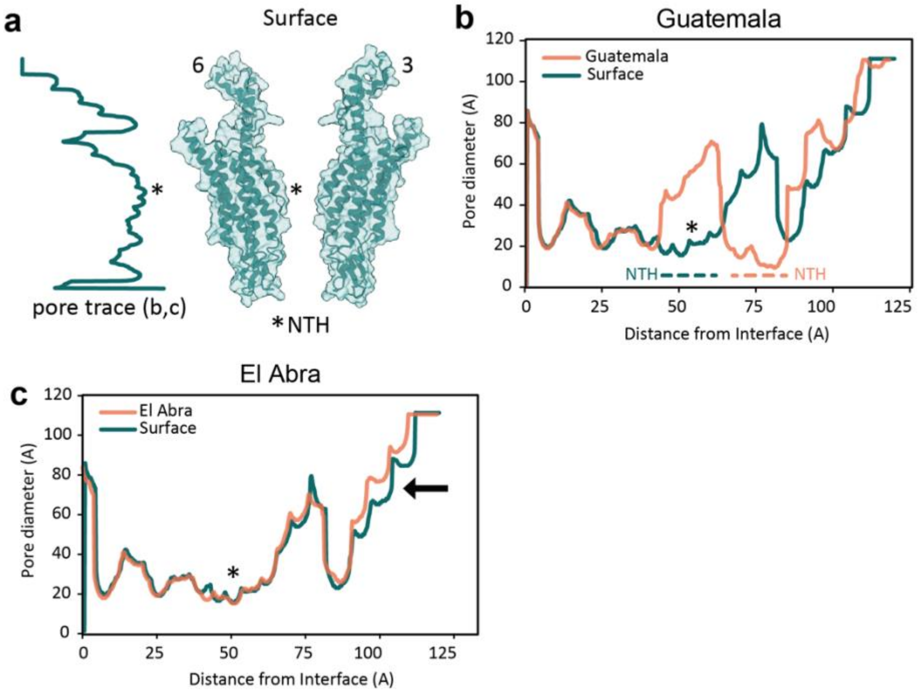

**Fig. S13. Traces of the topography of the interior of the Cx50 pore in *A. mexicanus*.** **(a)** Example of tracing the interior topography of the pore. Asterisk indicates the position of the downward surface NTH across all diagrams. **(b-c)** Pore traces are shown as pore diameter versus distance from the hemichannel-hemichannel interface. **(b)** Comparison of interior pore topography between the surface Cx50 and the Guatemala S89K mutation. The graphs show the NTH shift into the pore in the Guatemala lineage and resulting smaller pore diameter. **(c)** Comparison of the surface Cx50 and the El Abra T39M mutation. Pore traces are consistent between the surface and El Abra proteins except at the intracellular portion of the pore, at which point the El Abra pore is wider than the surface pore (arrow).

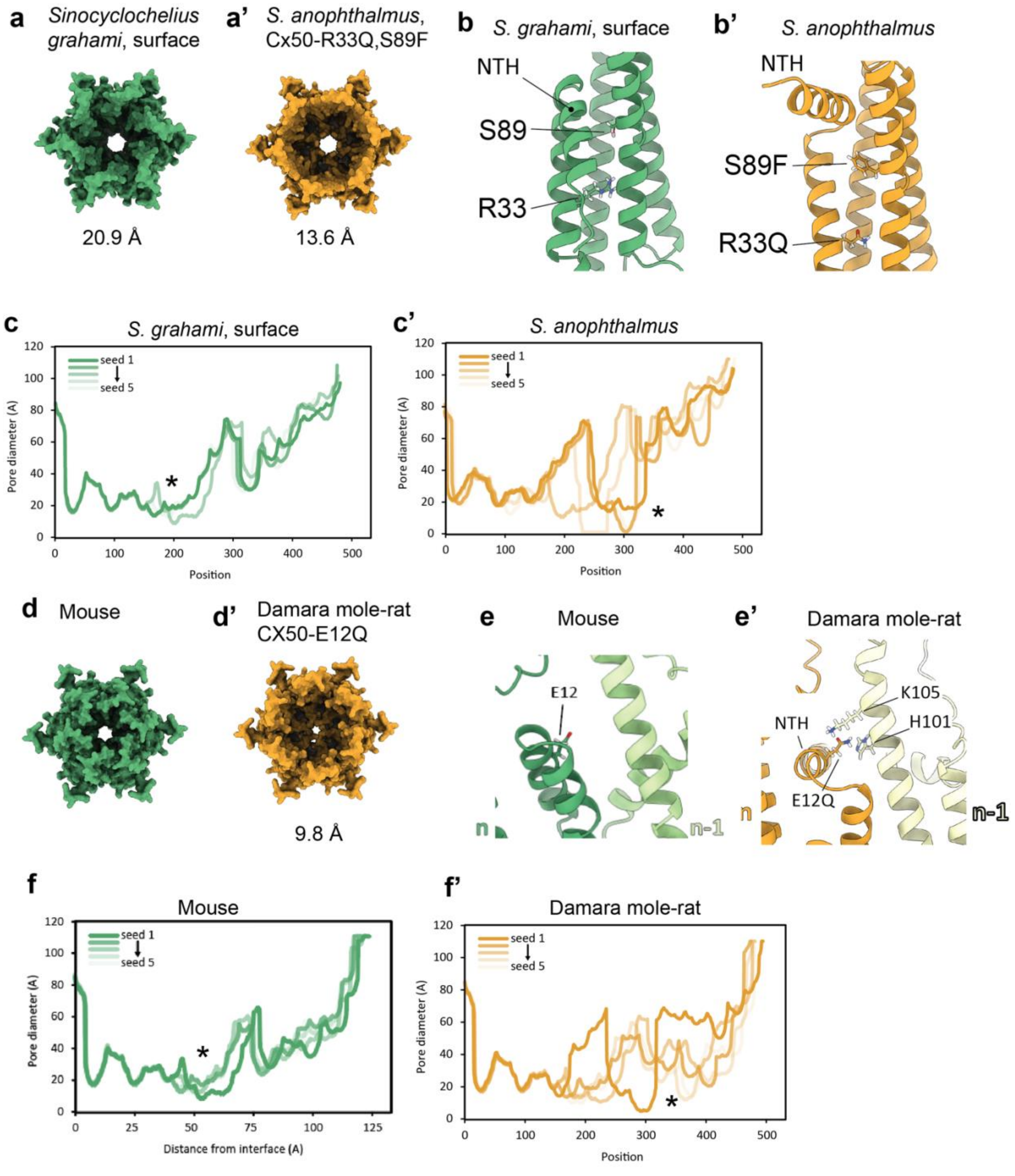

**Fig. S14. Predicted Cx50 and CX50 structures of select subterranean organisms.** **(a-c’)** Comparison Cx50 of the Chinese cavefish, *Sinocyclocheilus anophthalmus* and a close surface relative, *S. grahami*. **(a-a’)** *S. anophthalmus* has a smaller predicted pore size compared to the surface *S. grahami*. **(b-b’)** Close up modeling of S89 and R33 *Sinocyclocheilus* species. The S89F mutation is similar to the S89K mutation seen in the Guatemala lineage of *Astyanax mexicanus*, with phenylalanine occupying the space where the NTH is usually positioned close to transmembrane 2. In surface *S. grahami*,

R33 sits flush with adjacent helices in the same subunit. R33Q is slightly pushed out from the helix due to the several negatively charged residues that would normally interact with R33. **(c-c')** Traces of the interior of the pore show the frequent presence of the NTH into the pore in *S. anophthalmus* (asterisk) but not the surface fish *S. grahami* (asterisk). **(d-f')** Comparison of CX50 models in guinea pig and the subterranean Damara mole-rat, *Fukomys damarensis*. **(d-d')** The interior of the predicted pore is smaller in the Damara mole-rat compared to the model of CX50 from the mouse. **(e-e')** Close up of the E12 residue in mouse and Damara mole-rat. E12 in the mouse faces the interior of the pore as a hydrophobic residue. E12Q interacts with H101 in an adjacent subunit to stabilize the NTH in an upward configuration and orients away from the pore. **(f-f')** Interior pore tracing in the Damara mole-rat is highly variable with most traces showing the shift of the NTH position into the pore (asterisk) compared to traces in the mouse model (asterisk).

**Table S1. Population genomics summary statistics of the Caballo Moro cavefish.** Summary statistics were calculated using the popgenwindows.py package across overlapping 50kb windows (see Methods). The output includes absolute sequence divergence (Dxy) and relative sequence divergence (Fst) between pairwise populations and nucleotide diversity (pi) within each population.

**Table S2. SNPs and Indels that associate with eyeless CM fish but not eyed CM or surface fish.** Results of genome-wide association test - Cochran-Armitage trend test filtered to p<-log10(9) conducted with PLINK (*10*).

**Table S3. Eyeless specific SNPs with SnpEff annotation.** The SnpEff report of sites that were successfully annotated after genotype filtering. SNPs were filtered in the following manner: surface fish filtered to homozygous for the surface reference variant (RR), eyed Caballo Moro fish have at least one copy of the surface reference allele (RR or RA), eyeless Caballo Moro fish are homozygous for alternate alleles from the surface reference (AA). 203 variants met this criteria and were passed through SnpEff which predicts the effect of variants on protein function (predicted effect on coding genes and genomic locations; (*19*). Of the 203 variants provided to SnpEff, 89 were able to be annotated and fell within 40 genes, shown here. One gene has a predicted high impact effect on a protein coding region, *connexin50* or *gja8b*.

**Table S4. Candidate gene database.** A simple database of the genes identified as having potential involvement in eye loss. This database includes the genes identified by GWAS and genotype filtering in Table S2 and S3, as well as the genes identified from QTL analysis, a custom zebrafish gene ontology database, and a list of genes that have undergone hard selective sweeps in cave fish populations (see methods; *2*, *4*).

**Table S5. Subterranean and surface species used for Gja8b/GJA8 sequence comparison.** Sequences were pulled from NCBI. The closest surface relatives with available genomic sequences were identified with the NCBI Taxonomy Browser. * indicates reduced, but not absent eyes.

| Surface relative | Common name | Cavefish | Common name |
|---|---|---|---|
| *Astyanax mexicanus, surface* | Mexican tetra | *Astyanax mexicanus,* Guatemala | Mexican tetra, Guatemala lineage |
| | | *Astyanax mexicanus,* El Abra | Mexican tetra, El Abra lineage |
| *Ictalurus punctatus* | Channel catfish | *Rhamdia laluchensis* | La Luche cave catfish |
| | | *Rhamdia zongolicensis* | Zongolica catfish |
| | | *Trichomycterus rosablanca* | Andean cavefish |
| | | *Prietella phreatophila* | Mexican blindcat |
| | | *Trogloglanis pattersoni* | Toothless blindcat |
| *Danio rerio* | Zebrafish | | |
| *Cyprinus carpio* | Common carp | | |

| | | | |
|---|---|---|---|
| *Sinocyclocheilus grahami* | Golden-line barbel | *Sinocyclocheilus anshuiensis* | Blind barbel |
| *Sinocyclocheilus maitianheensis* | Golden-line barbel | *Sinocyclocheilus rhinocerous** | Horned golden-line barbel* |
| | | *Sinocyclocheilus anophthalmus* | Blind golden line fish |
| *Barbatula barbatula* | Stone loach | | |
| *Triplophysa tibetana* | Tibetan stone loach | *Triplophysa rosa* | Chinese blind loach |
| *Carapus acus* | Pearlfish | *Typhlias pearsei* | Mexican blind brotula |
| | | *Lucifuga dentata* | Toothed Cuban cusk-eel |
| *Percopsis omiscomaycus* | Trout-perch | *Forbesichthys agassizii* | Spring cavefish |
| | | *Forbesichthys papilliferus* | Northern spring cavefish |
| | | *Speoplatyrhinus poulsoni* | Alabama cavefish |
| | | *Amblyopsis rosae* | Ozark cavefish |
| | | *Amblyopsis hoosieri* | Hoosier cavefish |
| | | *Amblyopsis spelaea* | Northern cavefish |
| | | *Typhlichthys eigenmanni* | Eyeless cavefish |
| | | *Typhlichthys subterraneus* | Southern cavefish |
| **Surface relative** | **Common name** | **Subterranean mammal** | **Common name** |
| *Rattus norvegicus* | Brown rat | *Nannospalax galili* * | Upper Galilee blind mole-rat * |
| *Cavia porcellus* | Guinea pig | *Fukomys damarensis* * | Damara mole-rat * |
| | | *Heterocephalus glaber* * | Naked mole-rat * |
| *Erinaceus europaeus* | Western European hedgehog | *Talpa occidentalis* * | Iberian mole * |
| | | *Condylura cristata* * | Star-nosed mole * |

**Table S6. List of SRAs associated with PRJNA841601.**

### References

1. R. W. Mitchell, W. H. Russell, W. R. Elliott, Mexican Eyeless Characin Fishes, Genus Astyanax: Environment, Distribution, and Evolution. *Spec. Publ. Mus. Tex. Tech Univ.* (1977).

2. L. Espinasa, R. Borowsky, EYED CAVE FISH IN A KARST WINDOW. *J. Cave Karst Stud.* **62**, 180–183 (2000).

3. Z. Zhao, Y. Shen, Rain-induced weathering dissolution of limestone and implications for the soil sinking-rock outcrops emergence mechanism at the karst surface: a case study in southwestern China. *Carbonates Evaporites* **37**, 69 (2022).

4. L. Legendre, J. Rode, I. Germon, M. Pavie, C. Quiviger, M. Policarpo, J. Leclercq, S. Père, J. Fumey, C. Hyacinthe, P. Ornelas-García, L. Espinasa, S. Rétaux, D. Casane, Genetic identification and reiterated captures suggest that the Astyanax mexicanus El Pachón cavefish population is closed and declining. *Zool. Res.* **44**, 701–711 (2023).

5. Y. Yamamoto, L. Espinasa, D. W. Stock, W. R. Jeffery, Development and evolution of craniofacial patterning is mediated by eye-dependent and -independent processes in the cavefish Astyanax. *Evol. Dev.* **5**, 435–446 (2003).

6. R. J. Wassersug, A procedure for differential staining of cartilage and bone in whole formalin-fixed vertebrates. *Stain Technol.* **51**, 131–134 (1976).

7. Y. Elipot, H. Hinaux, J. Callebert, S. Rétaux, Evolutionary Shift from Fighting to Foraging in Blind Cavefish through Changes in the Serotonin Network. *Curr. Biol.* **23**, 1–10 (2013).

8. W. C. Warren, E. S. Rice, M. X, E. Roback, A. Keene, F. Martin, D. Ogeh, L. Haggerty, R. A. Carroll, S. McGaugh, N. Rohner, Astyanax mexicanus surface and cavefish chromosome-scale assemblies for trait variation discovery. *G3 GenesGenomesGenetics* **14**, jkae103 (2024).

9. M. A. DePristo, E. Banks, R. Poplin, K. V. Garimella, J. R. Maguire, C. Hartl, A. A. Philippakis, G. del Angel, M. A. Rivas, M. Hanna, A. McKenna, T. J. Fennell, A. M. Kernytsky, A. Y. Sivachenko, K. Cibulskis, S. B. Gabriel, D. Altshuler, M. J. Daly, A framework for variation discovery and genotyping using next-generation DNA sequencing data. *Nat. Genet.* **43**, 491–498 (2011).

10. S. Purcell, B. Neale, K. Todd-Brown, L. Thomas, M. A. R. Ferreira, D. Bender, J. Maller, P. Sklar, P. I. W. de Bakker, M. J. Daly, P. C. Sham, PLINK: A Tool Set for Whole-Genome Association and Population-Based Linkage Analyses. *Am. J. Hum. Genet.* **81**, 559–575 (2007).

11. R. L. Moran, E. J. Richards, C. P. Ornelas-García, J. B. Gross, A. Donny, J. Wiese, A. C. Keene, J. E. Kowalko, N. Rohner, S. E. McGaugh, Selection-driven

trait loss in independently evolved cavefish populations. *Nat. Commun.* **14**, 2557 (2023).

12. D. H. Alexander, J. Novembre, K. Lange, Fast model-based estimation of ancestry in unrelated individuals. *Genome Res.* **19**, 1655–1664 (2009).

13. J. S. Taylor, Y. Van de Peer, I. Braasch, A. Meyer, Comparative genomics provides evidence for an ancient genome duplication event in fish. *Philos. Trans. R. Soc. Lond. Ser. B* **356**, 1661–1679 (2001).

14. L. Excoffier, N. Marchi, D. A. Marques, R. Matthey-Doret, A. Gouy, V. C. Sousa, fastsimcoal2: demographic inference under complex evolutionary scenarios. *Bioinformatics* **37**, 4882–4885 (2021).

15. L. A. Bergeron, S. Besenbacher, J. Zheng, P. Li, M. F. Bertelsen, B. Quintard, J. I. Hoffman, Z. Li, J. St. Leger, C. Shao, J. Stiller, M. T. P. Gilbert, M. H. Schierup, G. Zhang, Evolution of the germline mutation rate across vertebrates. *Nature* **615**, 285–291 (2023).

16. N. Marchi, A. Kapopoulou, L. Excoffier, Demogenomic inference from spatially and temporally heterogeneous samples. *Mol. Ecol. Resour.* **24**, e13877 (2024).

17. N. Rosser, F. Seixas, L. M. Queste, B. Cama, R. Mori-Pezo, D. Kryvokhyzha, M. Nelson, R. Waite-Hudson, M. Goringe, M. Costa, M. Elias, C. Mendes Eleres de Figueiredo, A. V. L. Freitas, M. Joron, K. Kozak, G. Lamas, A. R. P. Martins, W. O. McMillan, J. Ready, N. Rueda-Muñoz, C. Salazar, P. Salazar, S. Schulz, L. T. Shirai, K. L. Silva-Brandão, J. Mallet, K. K. Dasmahapatra, Hybrid speciation driven by multilocus introgression of ecological traits. *Nature* **628**, 811–817 (2024).

18. E.-J. Wagenmakers, S. Farrell, AIC model selection using Akaike weights. *Psychon. Bull. Rev.* **11**, 192–196 (2004).

19. P. Cingolani, A. Platts, L. L. Wang, M. Coon, T. Nguyen, L. Wang, S. J. Land, X. Lu, D. M. Ruden, A program for annotating and predicting the effects of single nucleotide polymorphisms, SnpEff. *Fly (Austin)* **6**, 80–92 (2012).

20. J. Wiese, E. Richards, J. E. Kowalko, S. E. McGaugh, Quantitative trait loci concentrate in specific regions of the Mexican cavefish genome and reveal key candidate genes for cave-associated evolution. *J. Hered.*, esae040 (2025).

21. Y. Yamamoto, W. R. Jeffery, Central Role for the Lens in Cave Fish Eye Degeneration. *Science* **289**, 631–633 (2000).

22. Y. Yamamoto, W. R. Jeffery, Probing teleost eye development by lens transplantation. *Methods San Diego Calif* **28**, 420–426 (2002).

23. A. Dobin, C. A. Davis, F. Schlesinger, J. Drenkow, C. Zaleski, S. Jha, P. Batut, M. Chaisson, T. R. Gingeras, STAR: ultrafast universal RNA-seq aligner. *Bioinformatics* **29**, 15–21 (2013).

24. B. Li, C. N. Dewey, RSEM: accurate transcript quantification from RNA-Seq data with or without a reference genome. *BMC Bioinformatics* **12**, 323 (2011).

25. M. Labuhn, F. F. Adams, M. Ng, S. Knoess, A. Schambach, E. M. Charpentier, A. Schwarzer, J. L. Mateo, J.-H. Klusmann, D. Heckl, Refined sgRNA efficacy prediction improves large- and small-scale CRISPR-Cas9 applications. *Nucleic Acids Res.* **46**, 1375–1385 (2018).

26. E. M. Anderson, A. Haupt, J. A. Schiel, E. Chou, H. B. Machado, Ž. Strezoska, S. Lenger, S. McClelland, A. Birmingham, A. Vermeulen, A. van B. Smith, Systematic analysis of CRISPR–Cas9 mismatch tolerance reveals low levels of off-target activity. *J. Biotechnol.* **211**, 56–65 (2015).

27. B. A. Stahl, R. Peuß, B. McDole, A. Kenzior, J. B. Jaggard, K. Gaudenz, J. Krishnan, S. E. McGaugh, E. R. Duboue, A. C. Keene, N. Rohner, Stable transgenesis in Astyanax mexicanus using the Tol2 transposase system. *Dev. Dyn. Off. Publ. Am. Assoc. Anat.* **248**, 679–687 (2019).

28. J. Bibliowicz, A. Alié, L. Espinasa, M. Yoshizawa, M. Blin, H. Hinaux, L. Legendre, S. Père, S. Rétaux, Differences in chemosensory response between eyed and eyeless Astyanax mexicanus of the Rio Subterráneo cave. *EvoDevo* **4**, 25 (2013).

29. T. A. Hooven, Y. Yamamoto, W. R. Jeffery, Blind cavefish and heat shock protein chaperones: a novel role for hsp90alpha in lens apoptosis. *Int. J. Dev. Biol.* **48**, 731–738 (2004).

30. M. Mirdita, K. Schütze, Y. Moriwaki, L. Heo, S. Ovchinnikov, M. Steinegger, ColabFold: making protein folding accessible to all. *Nat. Methods* **19**, 679–682 (2022).

31. J. Jumper, R. Evans, A. Pritzel, T. Green, M. Figurnov, O. Ronneberger, K. Tunyasuvunakool, R. Bates, A. Žídek, A. Potapenko, A. Bridgland, C. Meyer, S. A. A. Kohl, A. J. Ballard, A. Cowie, B. Romera-Paredes, S. Nikolov, R. Jain, J. Adler, T. Back, S. Petersen, D. Reiman, E. Clancy, M. Zielinski, M. Steinegger, M. Pacholska, T. Berghammer, S. Bodenstein, D. Silver, O. Vinyals, A. W. Senior, K. Kavukcuoglu, P. Kohli, D. Hassabis, Highly accurate protein structure prediction with AlphaFold. *Nature* **596**, 583–589 (2021).

32. J.-J. Tong, U. Khan, B. G. Haddad, P. J. Minogue, E. C. Beyer, V. M. Berthoud, S. L. Reichow, L. Ebihara, Molecular mechanisms underlying enhanced hemichannel function of a cataract-associated Cx50 mutant. *Biophys. J.* **120**, 5644–5656 (2021).

33. E. F. Pettersen, T. D. Goddard, C. C. Huang, E. C. Meng, G. S. Couch, T. I. Croll, J. H. Morris, T. E. Ferrin, UCSF ChimeraX: Structure visualization for researchers, educators, and developers. *Protein Sci. Publ. Protein Soc.* **30**, 70–82 (2021).

34. W. Humphrey, A. Dalke, K. Schulten, VMD: Visual molecular dynamics. *J. Mol. Graph.* **14**, 33–38 (1996).

35. M. J. Abraham, T. Murtola, R. Schulz, S. Páll, J. C. Smith, B. Hess, E. Lindahl, GROMACS: High performance molecular simulations through multi-level parallelism from laptops to supercomputers. *SoftwareX* **1–2**, 19–25 (2015).

36. N. Michaud-Agrawal, E. J. Denning, T. B. Woolf, O. Beckstein, MDAnalysis: a toolkit for the analysis of molecular dynamics simulations. *J. Comput. Chem.* **32**, 2319–2327 (2011).

37. M. Stemmer, T. Thumberger, M. del Sol Keyer, J. Wittbrodt, J. L. Mateo, CCTop: An Intuitive, Flexible and Reliable CRISPR/Cas9 Target Prediction Tool. *PLoS ONE* **10**, e0124633 (2015).

38. J. P. Connelly, S. M. Pruett-Miller, CRIS.py: A Versatile and High-throughput Analysis Program for CRISPR-based Genome Editing. *Sci. Rep.* **9**, 4194 (2019).

39. J. Pang, N. Thomas, D. Tsuchiya, T. Parmely, D. Yan, T. Xie, Y. Wang, Step-by-step preparation of mouse eye sections for routine histology, immunofluorescence, and RNA in situ hybridization multiplexing. *STAR Protoc.* **2**, 100879 (2021).

40. W. R. Elliott, *The Astyanax Caves of Mexico: Cavefishes of Tamaulipas, San Luis Potosí, and Guerrero* (Association for Mexican Cave Studies, 2018).

41. J. Mazzoni, J. R. Smith, S. Shahriar, T. Cutforth, B. Ceja, D. Agalliu, The Wnt Inhibitor Apcdd1 Coordinates Vascular Remodeling and Barrier Maturation of Retinal Blood Vessels. *Neuron* **96**, 1055-1069.e6 (2017).

42. C. Zhang, Q. M. Turton, S. Mackinnon, K. K. Sulik, G. J. Cole, AGRIN FUNCTION ASSOCIATED WITH OCULAR DEVELOPMENT IS A TARGET OF ETHANOL EXPOSURE IN EMBRYONIC ZEBRAFISH. *Birt. Defects Res. A. Clin. Mol. Teratol.* **91**, 129 (2011).

43. G. Li, D. Jin, T. P. Zhong, Tubgcp3 Is Required for Retinal Progenitor Cell Proliferation During Zebrafish Development. *Front. Mol. Neurosci.* **12** (2019).

44. T. Li, Leber Congenital Amaurosis Caused by Mutations in RPGRIP1. *Cold Spring Harb. Perspect. Med.* **5**, a017384 (2015).
